# Microbial succession, community assembly and adaptation over five years in a newly discovered deep-sea cold seep

**DOI:** 10.1101/2024.10.31.619006

**Authors:** Qianyong Liang, Xinyue Liu, Jieni Wang, Tengkai Chen, Yingchun Han, Lu Zhang, Shuzhen Li, Jing Zhao, Yifei Dong, Binbin Guo, Xi Xiao, Xiyang Dong

## Abstract

Deep-sea cold seeps, characterized by the release of hydrocarbon-rich fluids such as methane from the seafloor, create unique ecosystems with evolving biological communities. Microorganisms serve as pioneers, initially colonizing early-stage cold seeps and providing essential matter and energy for subsequent colonizers. However, microbial succession, community assembly, and adaptation in these environments remain largely unexplored due to challenges associated with locating early-stage cold seeps. Here, we collected sediment samples from a newly discovered early-stage cold seep site, QDN-W07, in the South China Sea at varying depths over five years. We also sampled two previously reported cold seep sites, QDN-S18 and HM-ROV01, for comparative analysis. Amplicon sequencing revealed a high relative abundance of ANME-3 (up to 99%) and Methylomonadaceae (up to 83%) in surface sediments, and indicated changes in microbial community structures over time. Dispersal limitation dominated community assembly, while the contribution of homogeneous selection decreased over time, accompanied by a declined in the intensity of microbial association. Metagenomic analysis of core microbiome, comprising 172 metagenomic-assembled genomes at QDN-W07, revealed tolerance to low temperatures and oxygen. Core microbiome genomes were enriched with genes related to aerobic methane oxidation, urea metabolism, and the oxidation of sulfur, ammonia and carbon monoxide. These results indicate that core microorganisms obtain energy from the oxidation of inorganic compounds, allowing them to thrive in cold, methane-rich and unstable conditions. Overall, this study enhances our understanding of deep biosphere life, biogeochemical cycling, and the ecological management of cold seep environments.

## Introduction

Deep-sea cold seeps are typical marine habitats where hydrocarbon-rich fluids, primarily methane, naturally seep from the seafloor, creating localized environments rich in chemical energy (*1*). These fluids fuel unique ecosystems via chemosynthesis, influencing biogeochemical processes in sediments and composition of benthic communities (*2*). As cold seeps develop over time, community succession occurs, beginning with microbial colonization, followed by dense populations of clams or mussels, and then progressing to more complex and vibrant communities that include prevalent tubeworms and various macrofauna such as crabs, sea cucumbers and brittle stars (*3*, *4*). Active and mature cold seeps are relatively easy to discover due to their well-defined faunal communities and carbonate blocks (*2*), leading to extensive studies on community composition and succession processes in these ecosystems (*5*–*7*). In contrast, early-stage cold seeps are much harder to identify because they lack these well-established biological markers, and the dynamic flux of methane seeping from seabed complicates the discovery of new seep vents (*8*).

Microorganisms are pioneers in colonizing new cold seep sites (*9*, *10*), often forming thick mats on the seafloor. They utilize chemicals like methane, which is abundant in these environments, to generate energy and produce organic matter, serving as the cornerstone of cold seep ecosystems (*11*). For example, methane-oxidizing bacteria (MOB) and anaerobic methanotrophic archaea (ANME) obtain energy through aerobic and anaerobic methane oxidation (*12*) coupled with sulfate, nitrate, and nitrite reduction (*13*, *14*). These key microorganisms not only consume methane but also influence multiple elemental cycles and interact with other microorganisms (*15*). Additionally, these microbial activities lead to the formation of calcium carbonate on the seafloor, creating a stable substrate for later colonization by larger organisms (*9*, *16*, *17*). While microorganisms, especially methane-metabolizing microorganisms, play a foundational role in early-stage cold seeps, research on the composition and adaptation of microbial communities in this ecosystem remains limited.

Understanding the ecological forces that shape community assembly is essential for predicting microbial functions and biodiversity patterns in microbial ecology (*18*). According to Vellend’s theory (*19*), community assembly patterns are influenced together by four key processes: selection, drift, speciation, and dispersal. In mature cold seep sediments, an increased supply of methane could lead to stronger homogeneous selection, favoring the dominance of methanotrophic and sulfate-reducing microorganisms (*5*). In early-stage cold seeps, where methane begins to emerge, other environmental factors, such as carbonates and metal ions, also change (*9*). Dispersal is critical in environments with dramatic shifts, enabling microorganisms to colonize new niches and adapt to changing conditions (*18*). Despite these insights, it remains unclear which community assembly processes drive microbial community changes in early-stage cold seeps and which specific factors influence this assembly.

In this study, we collected sediment samples from the newly discovered cold seep site QDN-W07, located approximately 50 km northeast of the reported Haima cold seep (*20*) in the Qiongdongnan Basin (QDNB) of the South China Sea **(Fig. 1A)**. Sampling was conducted over five-year, from 2019 to 2023, resulting in a total of 184 samples obtained from the seepage and its surrounding area within a radius of 30 to 500 m. The sediment cores reached a maximum depth of 720 centimeters below the seafloor (cmbsf) **(Table S1)**. For comparative purposes, we also collected 48 samples from two previously reported seep sites, HM-ROV01 and QDN-S18, within the Haima cold seep. Our research objectives were threefold: (1) to outline succession patterns of cold seep microbial communities over time and space; (2) to investigate the abiotic and biotic factors that influence the assembly and progression of these microbial communities; and (3) to delineate the composition, adaptation and metabolic potential of core microbiome at QDN-W07. Through this investigation, we aim to enhance understanding of microbial community assembly processes, the role of core microbiome, and ecological mechanisms underpinning methane metabolism in the distinctive environment of cold seeps.

**Figure 1.**
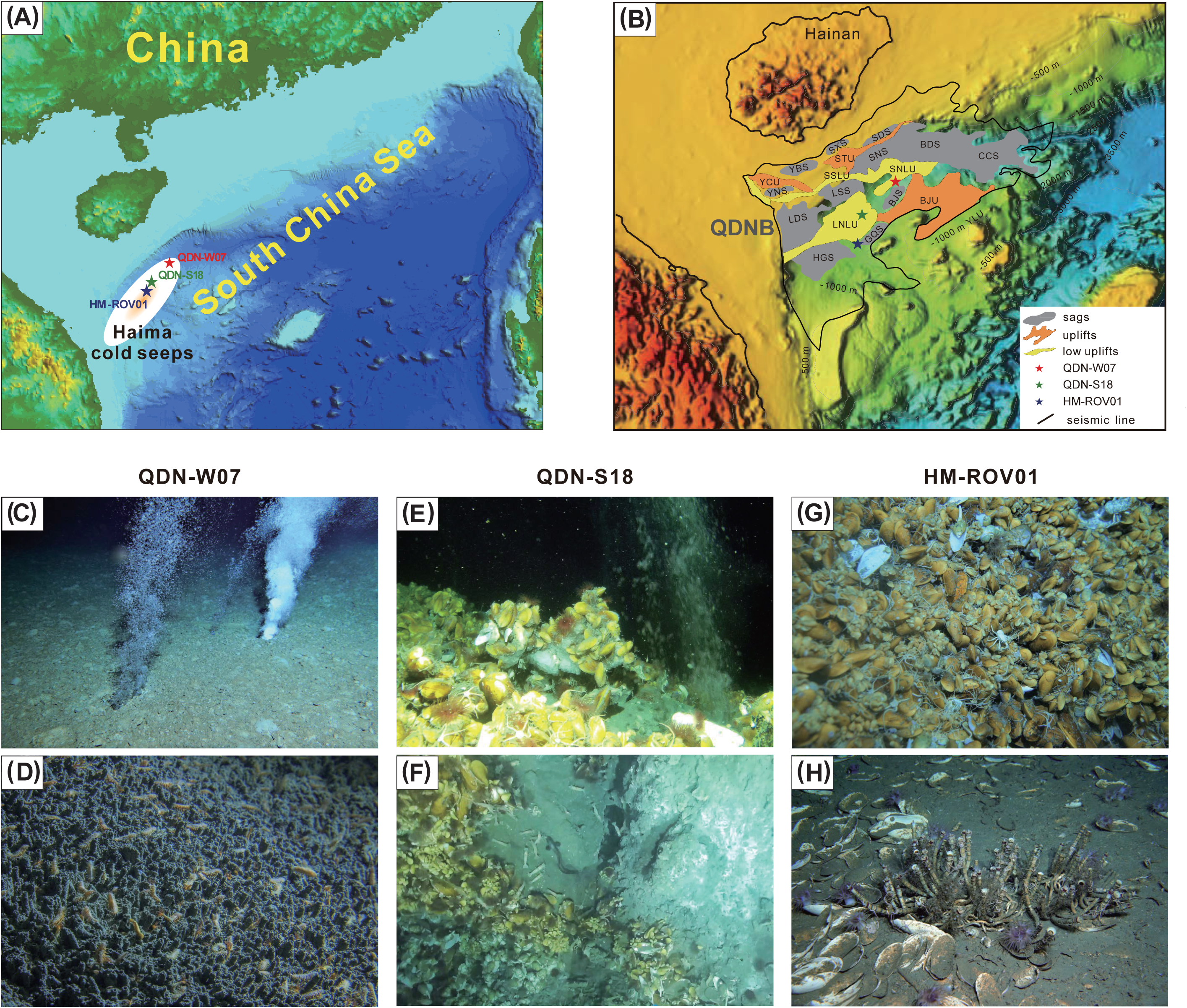
Geographical and ecological overview of three seep sites. **(A)** Bathymetry map of the study area. **(B)** Geological settings of three cold seep sites in the Qiongdongnan Basin (QDNB), including HM-ROV01 and QDN-S18, and newly discovered QDN-W07. **(C)** Methane seepage and **(D)** polychaeta benthos at QDN-W07. **(E)** Mussels living with methane hydrates and bubbles and **(F)** carbonate crusts at QDN-S18. **(G)** Dense mussel beds and **(H)** tube worms with dead clam shell debris at HM-ROV01. BJS: Beijiao Sag. BDS: Baodao Sag. CCS: Changchang Sag. GQS: Ganquan Sag. HGS: Huaguang Sag. LDS: Ledong Sag. LSS: Lingshui Sag. SDS: Songdong Sag. SNS: Songnan Sag. SXS: Songxi Sag. YBS: Yabei Sag. YNS: Yanan Sag. BJU: Beijiao Uplift. STU: Songtao Uplift. YCU: Yacheng Uplift. YLU: Yongle Uplift. LNLU: Lingnan Low Uplift. LSLU: Low Uplift. SNLU: Songnan Low Uplift.

## Results and discussion

### Newly discovered cold seep QDN-W07 was identified as an early-stage cold seep

The research area is located in the QDNB with a structural framework of three depressions and two uplifts (*20*) on the lower continental slope of the northwestern South China Sea (**Fig. 1, A and B**). The newly discovered cold seep, named QDN-W07, lies in the Songnan Low uplift at a water depth of 1,766 m and is connected to a deep gas chimney beneath the seafloor (**Fig. 1B**). The gas chimney originates from a basal uplift, appearing as a blanking zone in seismic sections (**Fig. S1A**) with an approximate width of 1 km. Above this zone, enhanced reflections are visible, and the gas chimney narrowed upwards to less than 200 m, appearing as columnar, vertically convex-aligned reflectors. Ultimately, it manifests as a crater with a diameter of 300 m on the seabed (**Fig. S1B**). Analysis from the same site GMGS5-W07, drilled during the fifth “China National Gas Hydrate Drilling Expedition” (GMGS5) in 2018, indicates that the system had experienced dynamic changes in fluid flux and multi-stage gas hydrate evolution over a time scale ranging from months to thousands of years (*21*).

The development stage of the QDN-W07 site was assessed based on habitat characteristics (*9*, *22*, *23*), including rapid methane emission and extensive bacterial mats (**Fig. 1C**), along with a high abundance of polychaetes (e.g., *Spionidae*) on the seafloor, without aggregations of large fauna (**Fig. 1D**). These features indicate that QDN-W07 represents an early stage of cold seep ecosystem development (*24*). In contrast, the reported active and mature cold seep sites, HM-ROV01 (*23*) and QDN-S18 (*20*) (**Fig. 1B**), exhibit more complex communities. The QDN-S18 site, characterized by high fluid flux, supports dense mussel beds, sea cucumbers, anemones and massive carbonate deposition **(Fig. 1, E and F)**. Meanwhile, the HM-ROV01 site, which experiences lower methane seepage intensity, hosts a diverse range of organisms, including living mussels (*Bathmodiolus plantifrons*), peripheral tube worms (*Paraescarpia echinospica*) and dead clams (mainly *Calyptogena* sp.) **(Fig. 1, G and H).**

### Geochemical data indicate progressive methane oxidation at QDN-W07

Geochemical analysis of piston core samples collected from the QDN-W07 site over multiple years (2019, 2022, and 2023) reveals distinct depth profiles compared to those from the QDN-S18 site in 2022 (**Fig. 2 and Table S2**). Sedimentary CH_4_ concentrations across all four cores remained high (153∼1,795 μM) and fluctuated violently (**Fig. 2A**). At both the QDN-W07 site in 2022 and 2023, and at the QDN-S18 site, CH [concentration patterns were similar, with peak values occurring between 0 and 200 cmbsf, followed by subsequent decreases. Elevated surface sediment CH [levels, along with visually observed methane seepage, suggest that methane is venting directly from deeper sediments to the seafloor. In core QDN-W07-2019, dissolved inorganic carbon (DIC) concentrations and stable carbon isotope ratios of DIC (δ^13^C_DIC_) fluctuated slightly, ranging from 1.5 to 2.7 mM and -2.5 to 0.5‰ (**Fig. 2, B and C**), aligning closely with typical seawater values (*25*). In core QDN-W07-2022, DIC remained stable, while δ^13^C_DIC_ initially rose sharply (from -25‰ to - 6‰) before stabilizing, indicating that initial microbial degradation of organic matter or methanogenesis occurred above 220 cmbsf. Core QDN-W07-2023 exhibited a pattern similar to that of QDN-S18, indicative of typical organic matter degradation and methane oxidation, characterized by increasing DIC levels and more negative δ^13^C_DIC_ values; however, it was less reactive. These findings suggest that over the five-year period at QDN-W07, microbial-mediated methane oxidation began at the sediment surface and gradually extended to deeper layers (*26*).

**Figure 2.**
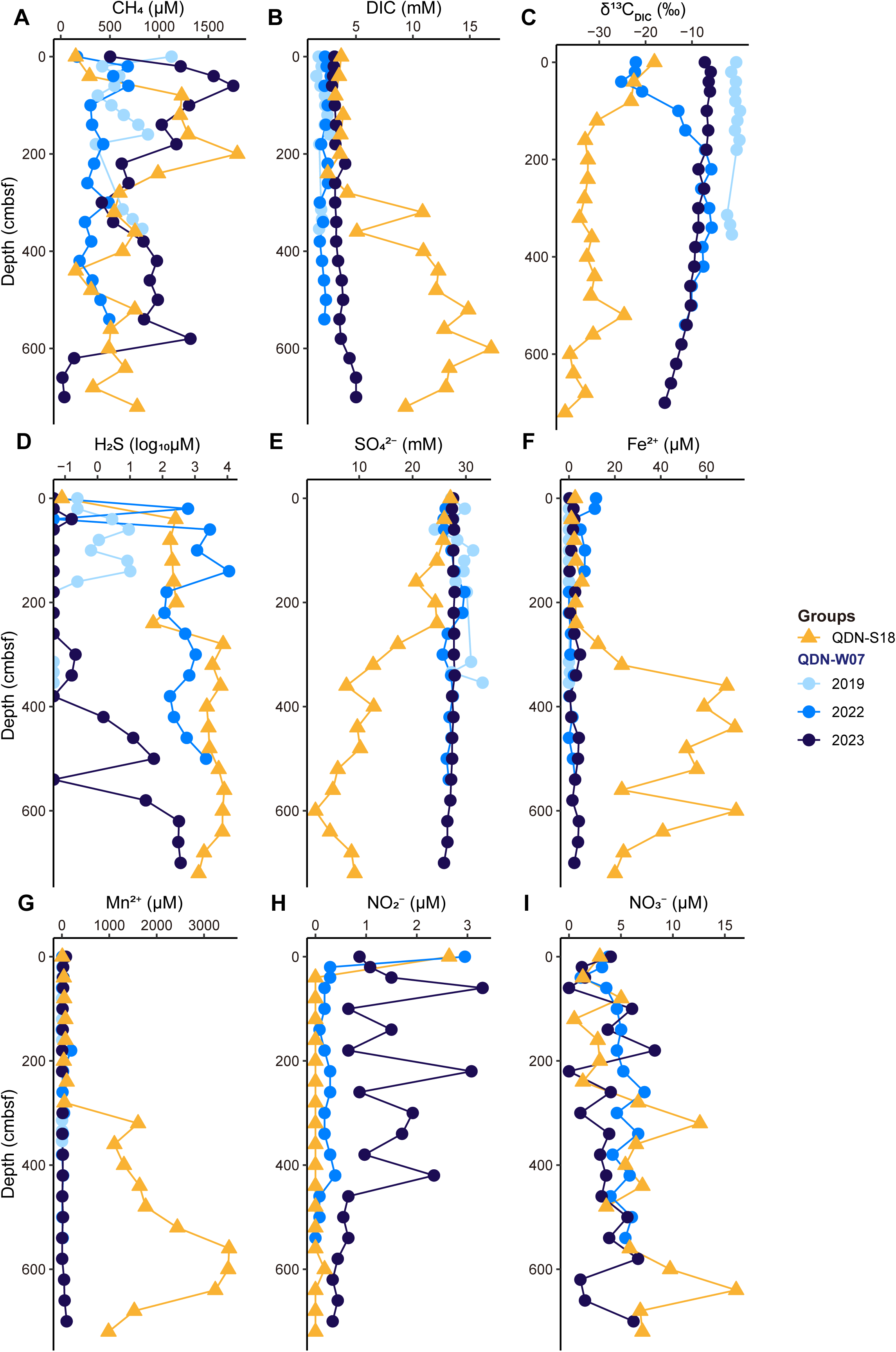
Depth profiles of geochemical parameters at QDN-W07 in 2019, 2022, 2023 and QDN-S18 cold seeps. The plots present the concentration gradients of various chemicals measured at different sediment depths, including **(A)** methane (CH_4_), **(B)** dissolved inorganic carbon (DIC), **(C)** carbon isotopic composition of dissolved inorganic carbon (δ^13^C_DIC_), **(D)** hydrogen sulfide (H_2_S), **(E)** sulfate (SO_4_^2-^), **(F)** ferrous ion (Fe^2+^), **(G)** manganese ion (Mn^2+^), **(H)** nitrite (NO_2_ ^-^), and **(I)** nitrate (NO_3_^-^). Source data are provided in **Table S2**.

Hydrogen sulfide (H_2_S) levels were comparatively lower at QDN-W07 in 2019 and 2023 (up to 313 μM and 363 μM) compared to those at QDN-W07 in 2022 and QDN-S18 (up to 11,244 μM and 7,967 μM) **(Fig. 2D)**. Sulfate (SO[²[) concentrations remained relatively high (near to the average value in seawater) (*25*) and stable at QDN-W07, while they declined markedly at QDN-S18 (**Fig. 2E**). Other factors, such as dissolved ferrous iron (Fe²[) and manganese (Mn²[), were more stable and lower across different depths at QDN-W07 than at QDN-S18 (**Fig. 2, F and G**). These results indicate that, anaerobic oxidation of methane (AOM) is more intense at the active cold seep QDN-S18, likely coupled with sulfate or metals reduction (*9*, *27*). Interestingly, nitrite (NO[[) variations were limited to above the depth of 420 cmbsf in QDN-W07-2023 and were nearly undetectable in QDN-S18 (**Fig. 2H**). Nitrate (NO_3_^-^) concentrations varied slightly in all piston cores (**Fig. 2I**). These observed geochemical differences likely stem from varied geological activity and depth-dependent shifts in microbial biogeochemical processes.

### Methylomonadaceae and ANME-3 are abundant at QDN-W07

Based on 16S rRNA gene amplicons, which yielded 47,676 archaeal amplicon sequence variants (ASVs) and 82,084 bacterial ASVs, microbial community compositions at QDN-W07 exhibited obvious variations across sites, depth and year (**Figs. 3 and 4, Tables S3 and S4**). Alpha-diversity analysis using the Shannon index showed that the microbial community at QDN-W07 was more diverse than these at active cold seeps, particularly among archaea (**Fig. S2, A and B,** Wilcoxon test, *p* < 0.0001), highlighting the unique microbial characteristics of QDN-W07 as an early-stage cold seep. At QDN-W07, both archaeal (ANOSIM, *R* = 0.29, *p* = 0.001) and bacterial (*R* = 0.44, *p* = 0.001) community structures varied significantly with depth (**Fig. S3, A and B**). This depth-related differentiation was further supported by a clear depth-decay pattern, especially for bacterial communities (*R^2^*> 0.1, *p* < 0.001) (**Fig. S4**), indicating community similarity decreased with increasing depth (*28*, *29*). Specifically, surface sediments were dominated by archaeal phylum Halobacrerota and bacterial phylum Proteobacteria, while deeper sediments showed an obvious increase in the relative abundance of archaeal phyla Asgardarchaeota and Crenarchaeota, and bacterial phylum Chloroflexi (**Fig. 3, A and B**). Moreover, archaeal (PERMANOVA, *R^2^* = 0.12, *p* < 0.001) and bacterial (*R^2^* = 0.07, *p* < 0.001) community compositions in seepage areas showed only weak differences from those in surrounding areas at QDN-W07, suggesting that microbial communities at QDN-W07 still share some similarities with those in non-seep sediments. Over the five -year study period, although the predominant phyla at QDN-W07 remained consistent from 2019 to 2023, their relative abundances fluctuated, revealing interannual variations in the sediment microbial community structure.

**Figure 3.**
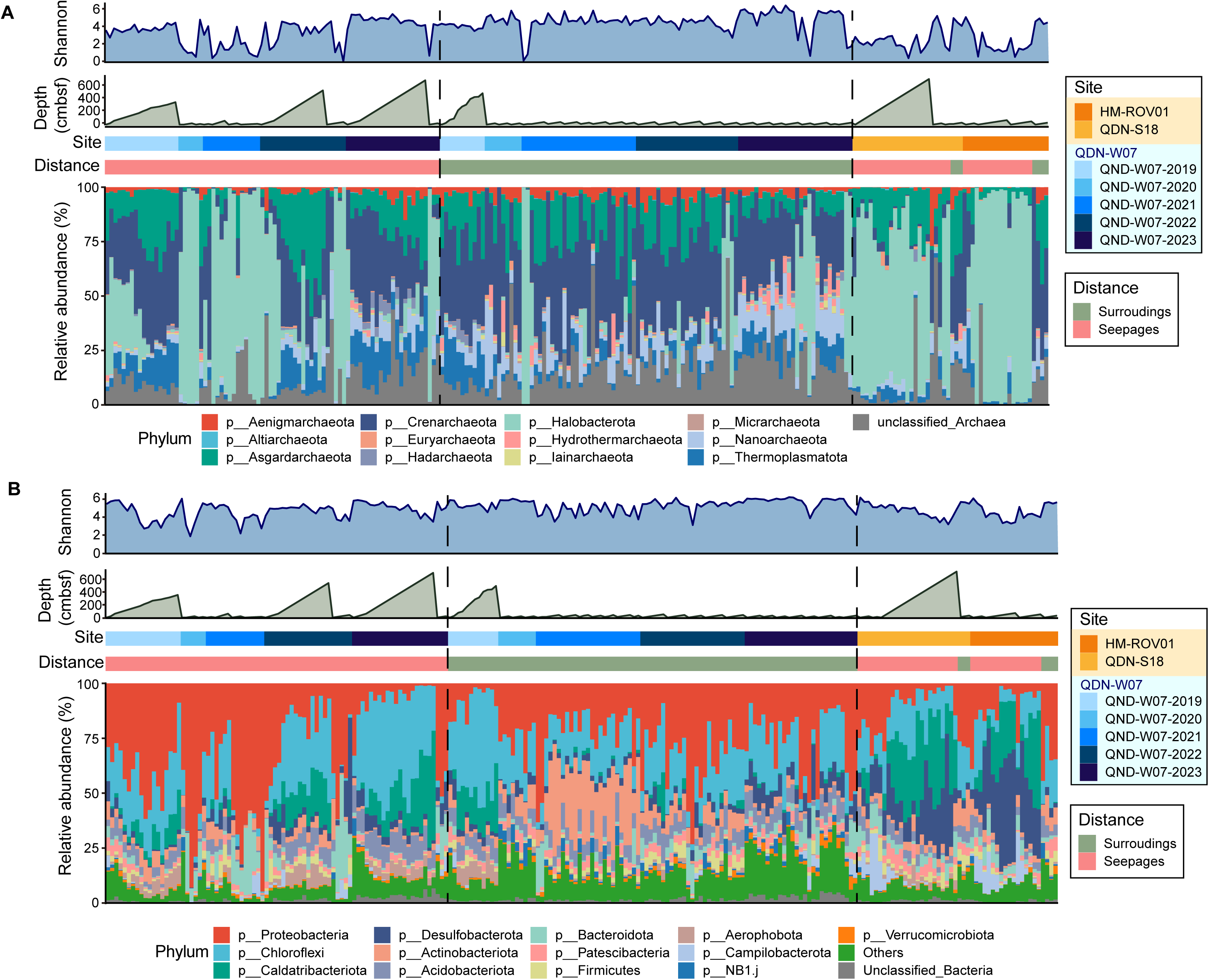
Taxonomic composition of microbial communities at three cold seep sites based on 16S rRNA gene amplicon. The bar charts show relative abundance of archaeal phyla **(A)** and bacterial phyla **(B)** across different samples. In both panels, the upper blue line charts show the Shannon index of corresponding domain, while the green line charts indicate sediment depth for each sample. The color bars above the bar charts indicate the sample source, including the site, sampling year and distance from seep (seepage and surrounding area). Source data are provided in **Table S3**.

**Figure 4.**
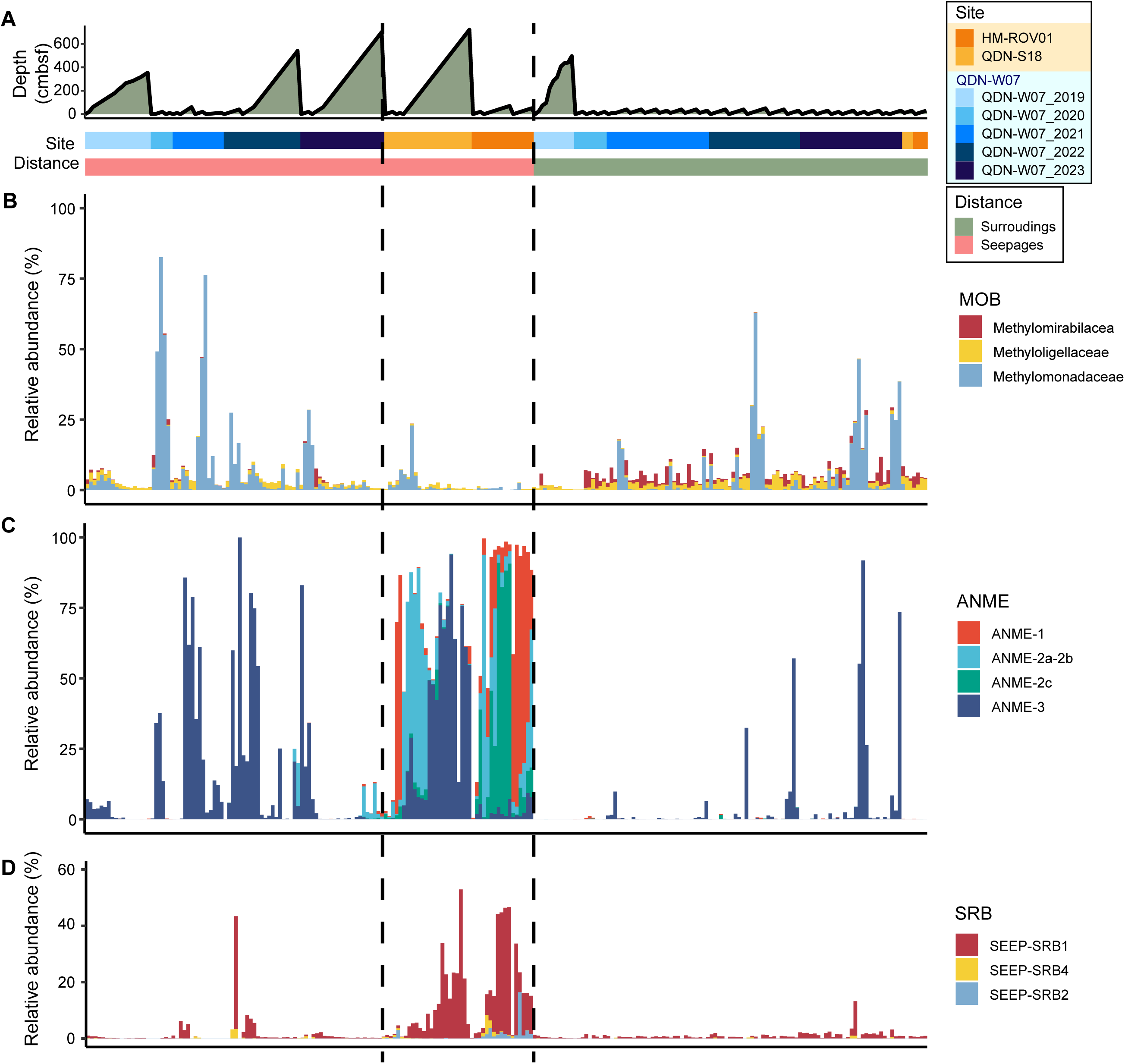
Relative abundance of important microbial taxa across different years, depths, sites, and distances from seepage areas. **(A)** Sources and sediment depth of samples. The green line charts show sediment depth of each sample, while the color bars below indicate the sample source, including the site, sampling year and distance from seep (seepage and surrounding area). Relative abundance (%) of **(B)** methane-oxidating bacteria (MOB), **(C)** anaerobic methanotrophic archaea (ANME) and **(D)** sulfate-reducing bacteria (SRB) across different samples. Source data are provided in **Table S4**.

The QDN-W07 site featured abundant methane-metabolizing microorganisms, particularly Methylomonadaceae and ANME-3 (**Fig. 4**). In surface sediments, aerobic methane-oxidizing bacteria Methylomonadaceae were notably prevalent, especially in 2020 and 2021, while their presence decreased in deeper layers. In contrast, Methylomonadaceae were less abundant at the QDN-S18 and HM-ROV01 sites (**Fig. 4B, Table S4**). Our data revealed a substantial presence of ANME within seepage area, while they were less prevalent in surrounding area **(Fig. 4C, Fig. S5 and Table S4)**. Both QDN-W07 and QDN-S18 exhibited relatively high abundances of ANME-3, similar to previous observations in other cold seeps like the Haakon Mosby mud volcano (*15*, *30*, *31*). However, a notable difference was that at QDN-W07, ANME-3 was primarily distributed in the surface and shallow layers, with virtually no other ANME groups present, while at QDN-S18, it was mainly found in deeper layers. Furthermore, HM-ROV01 and QDN-S18 exhibited more ANME, especially ANME-1 and ANME-2, whereas QDN-W07 sediments exhibited lower ANME abundances with lower sulfate-reducing bacteria (SRB) abundances **(Fig. 4D)**. Over five years at QDN-W07, methane-metabolizing microorganisms were scarce in 2019, while the relative abundance of Methylomonadaceae peaked in 2020 and then declined, and ANME-3 increased steadily from 2020 to 2023. This trend indicates that methane oxidation processes at the early-stage cold seep QDN-W07 are strengthening over time, shifting from aerobic methane oxidation (AeOM) to AOM, and suggesting that ANME microorganisms may become more diverse and expand from the surface to deeper sediments in the future.

### Microbial interactions decline and environmental influences increase during cold seep development

The interrelationships among microbial taxa and their associations with environmental variables drove the variations in microbial community structure over time at QDN-W07. To explore microbial interactions, such as competition and mutualism (*32*, *33*), co-occurrence networks of microorganisms in seepage areas were constructed **(Fig. 5A)**, excluding QDN-W07-2020 due to limited samples (*34*). The results showed a predominance of intra-domain connections over inter-domain connections, with most correlations being positive. Such patterns typically enhance network stability and resistance to disturbances (*35*). Time series analysis revealed that as cold seep developed, the average degree of networks decreased **(Fig. 5B**, *R^2^* = 0.48, *p* = 0.078), particularly within the bacterial community **(Fig. 5A and Table S5)**, indicating a decline in microbial interactions over time. This trend was also observed in the relationship between bacteria and archaea, as both the number of nodes and the average degree of inter-domain connections declined **(Fig. S6)**. Notably, the proportion of negative correlations, often indicative of competition or niche differentiation (*36*), decreased from 30.6% in 2019 to 5.6% in 2023 (**Table S5)**. These above findings suggest that co-exclusion mechanisms, such as competition, were more common among microbial communities at early-stage cold seeps. As development progressed, niche differentiation among microorganisms may have occurred (*37*), leading to reduced diversity and fewer negative correlations. This is consistent with lower alpha diversity observed at two active cold seeps compared to QDN-W07 **(Fig. S2)**. Consequently, the surviving species, selected by the extreme conditions of cold seeps, exhibited more facilitative interactions with one another.

**Figure 5.**
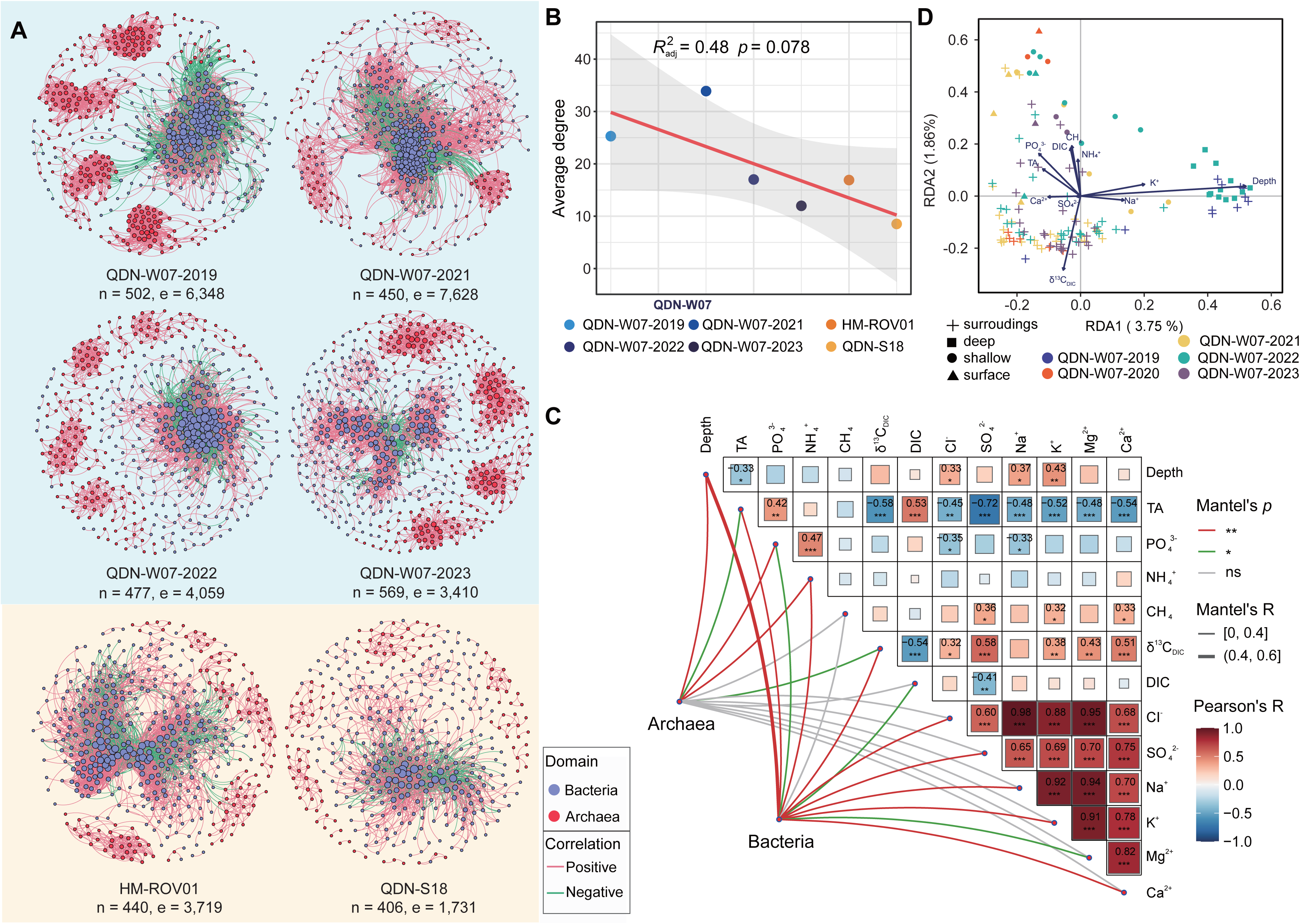
Microbial co-occurrence networks and environmental drivers in cold seep sediments of three sites. **(A)** Microbial co-occurrence networks of microbes sampled from different sites and during different years. Each network is characterized by the number of nodes (n), representing species, and the number of edges (e). Nodes are colored by domain, with red nodes representing archaea and blue nodes representing bacteria. Node size indicates the degree of connectivity, while red edges represent significant positive correlations, whereas green edges represent significant negative correlations. **(B)** Scatter plot shows the correlation between average degree of co-occurrence networks and different groups. Source data of **(A)** and **(B)** are provided in **Table S5**. **(C)** Distance-based redundancy analysis (dbRDA) of microbial community composition in relation to environmental gradients at QDN-W07. **(D)** Mantel correlation analysis between geochemical data and microbial communities at QDN-W07. Edge width corresponds to Mantel R statistic, while edge color denotes the statistical significance. Pairwise comparisons of geochemical data are shown with a color gradient denoting Pearson’s correlation. Source data for panels **(C)** and **(D)** are provided in **Table S6**.

At QDN-W07, microbial communities displayed minimal correlation with environmental variables, being substantially impacted only by depth (Mantel test, *R* = 0.44, *p* = 0.001) **(Fig. 5C and Table S6)**. In contrast, bacterial communities at the more active sites HM-ROV01 and QDN-S18, were influenced by several environmental factors, including SO_4_^2-^, Ca^2+^, and depth **(Fig. S7, A and B and Table S6)**. The distance-based redundancy analysis (dbRDA) **(Fig. 5D)** further demonstrated clear depth stratification; surface and shallow microorganisms correlated strongly with CH_4_, DIC and PO_4_ ^3-^, while δ^13^C mainly impacted microorganisms in surrounding areas. However, the low explanatory power of each constrained axis (< 4%, **Fig. 5C**) indicated that environmental factors could not adequately explain the distribution of microbial community composition at QDN-W07, aligning with the Mantel test results. Interestingly, analyses over the five-year period indicated a trend of increasing correlation with environmental factors and decreasing inter-species interactions as cold seeps developed.

### Dispersal limitation and homogeneous selection were dominant community assembly processes at QDN-W07

The quantitative analysis of microbial community assembly processes (*38*) showed that across all sites, deterministic process of homogeneous selection (HoS) and stochastic process of dispersal limitation (DL) dominated, shaping over 95% of community structures (**Fig. 6, A and B, and Table S7**). Archaeal communities at QDN-W07, compared to these at active cold seeps, were more influenced by HoS (33.3%-70.0%, **Fig. 6A and Fig. S8C**), suggesting higher environmental stress on archaea (*39*). Over five years, the influence of HoS at QDN-W07 consistently decreased, although it remained still higher than at HM-ROV01 and QDN-S18 (*R^2^* = 0.63, *p*= 0.037; **Fig. 6C and Fig. S8C**). Microbial interactions, such as competition, can create environmental stress on microorganisms, driving selection processes (*33*, *40*). Therefore, the reduction in microbial interactions (**Fig. 5A**) may contribute to the decrease in HoS at QDN-W07. Since HoS typically increases community similarity (*39*), the microbial community at QDN-W07 shifted from a more diverse assemblage to a less diverse, ANME-dominated one (**Fig. 4 and Fig. S2**). At HM-ROV01 and QDN-S18, the dominance of ANME (**Fig. 4**) suggests that these communities were more adapted to cold seep conditions, facing lower selective pressures and experiencing weaker HoS effects (**Fig, 6A**). Spatially, HoS had the greatest impact on surface-layer archaeal communities at QDN-W07, with its influence diminishing with depth. Conversely, the opposite trend was observed at the other two sites (**Fig. 6D and Fig. S8, A to C**).

**Figure 6.**
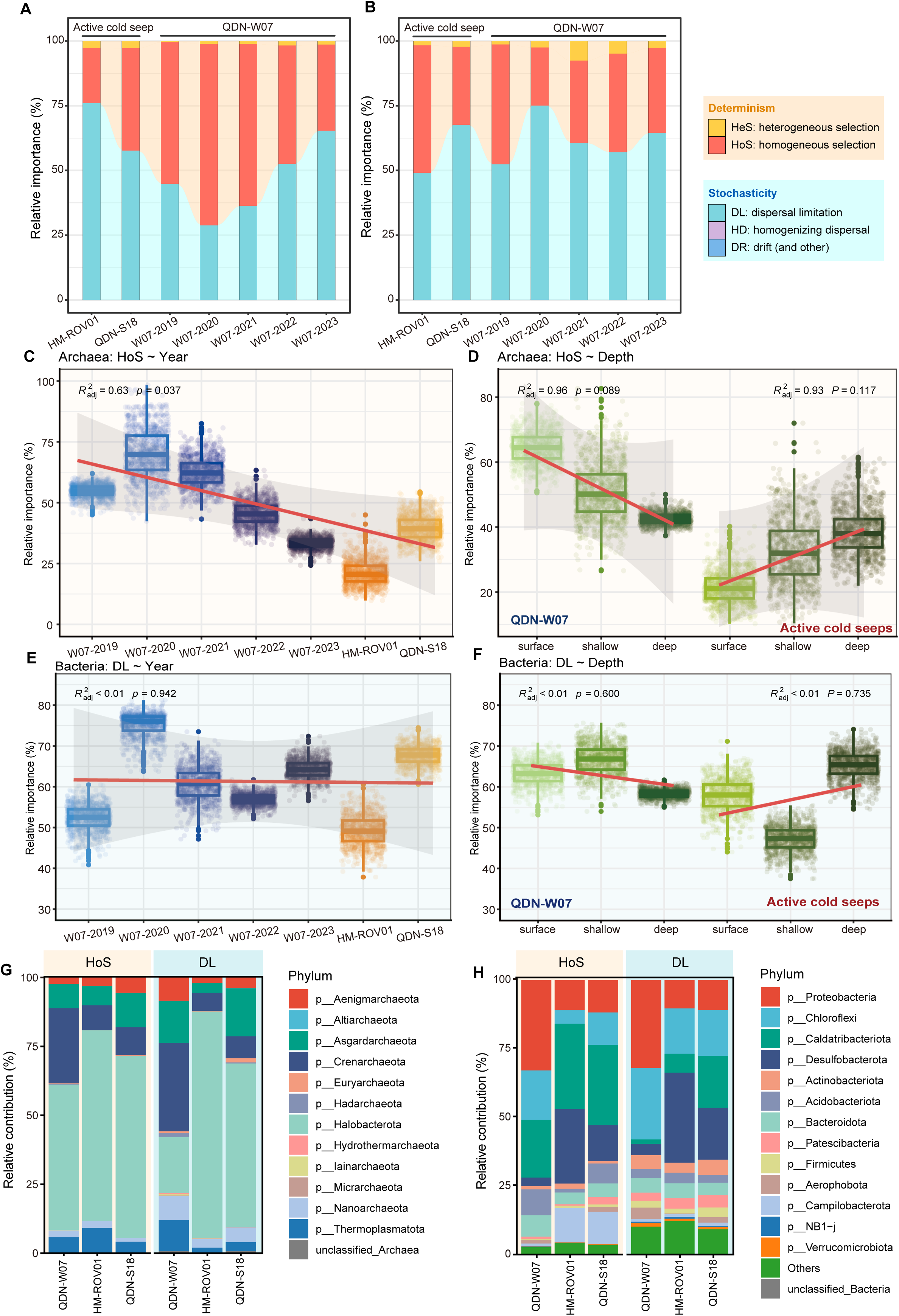
Microbial community assembly processes at QDN-W07, HM-ROV01 and QDN-S18. Relative contributions of five ecological processes for **(A)** archaeal and **(B)** bacterial communities across three cold seeps. Relative importance of **(C-D)** homogeneous selection (HoS) for archaeal and **(E-F)** dispersal limitation (DL) for bacterial communities at different sites and sediment depths. The boxplots are based on bootstrapping results (1,000 times). Red line indicates trendline from linear regression, with grey shaded area representing 95% confidence interval. *R^2^*and *p* values show the coefficient of determination and significance of the regression, respectively. In **(C)** and **(E)**, blue boxes indicate data from QDN-W07, with color intensity reflecting variation across different years. **(D)** and **(F)** display the trends in **(D)** HoS for archaea and **(F)** DL for bacteria across sediment depths. The left side of them shows the trends at QDN-W07, and the right side shows the trends at active seeps, including HM-ROV01 and QDN-W07. Relative contributions of different **(G)** archaeal and **(H)** bacterial phyla to HoS or DL at different sites. In each panel, the left side (orange) represents HoS, and the right side (blue) represents DL. Source data are provided in **Tables S7-10**.

Overall, DL predominantly controlled microbial community assembly, particularly for bacterial communities (**Fig. 6, A and B)**, with no significant trends over time or depth **(Fig. 6, E and F, Fig. S8C)**. In non-seep marine sediments, microbial dispersal is largely passive and severely limited, with microorganisms taking around a million years to move just 6 m vertically (*41*). In cold seeps, upwelling fluids from deeper sediments facilitate the passive dispersal of deep biosphere microorganisms from subsurface to surface environments (*42*). At QDN-W07, the lack of extensive carbonate precipitation and macrofaunal coverage **(Fig. 1C)** potentially enhances upwelling fluid flow. This may explain why DL has a lesser effect on archaea and deep-layer bacteria at QDN-W07 compared to the other two active cold seeps **(Fig. 6F)**. Furthermore, DL contributes to great divergence in community structure (*40*), as evidenced by a clear depth-decay pattern where community similarity notably decreases with increasing depth, especially for bacteria (*p* < 0.001, **Fig. S4)**.

The temporal and spatial dynamics of microbial community assembly at QDN-W07 are influenced by both environmental factors and microbial composition. In 2020, the strongest associations between HoS and environmental factors were observed. Archaeal communities were negatively influenced by the difference of depth and Ca^2+^ concentrations, while bacterial communities were positively associated with CH_4_ concentrations **(Fig. S9 and Table S8)**. At QDN-W07, predominant ‘bins’ with high abundant are more affected by HoS. In details, merely 3.7% of archaeal ‘bins’ (phylogenetic groups) and 3.2% of bacterial ‘bins’ were were primarily affected by, yet their relative abundance up to 24.6% and 26.2%, respectivel. In contrast, relatively rarer ‘bins’ are more susceptible to DL **(Table S9 and S10)**. These results were consistent with microbial community assembly patterns observed in other environments (*43–46*).

Specifically, highly abundant taxa such as archaeal phylum Halobacterota **(Fig. 3A)** and bacterial phylum Caldatribacteriota **(Fig. 3B)** obviously contributed less to DL than to HoS **(Fig. 6, G and H)**. In contrast, rarer taxa, such as Nanoarchaeota and Actinobacteriota, were more influential in DL **(Fig. 6, G and H)**. Abundant taxa tend to possess wider niches, allowing them to utilize a broader range of resources and disperse more easily, thus experiencing less DL (*45*).

### Core microbiome at QDN-W07 exhibits greater tolerance to low temperature and oxygen

Core microbiome represents a stable and persistent microbial component within the habitat, providing a relatively “invariant” baseline amidst dynamically changing communities (*47*, *48*). Eighteen archaeal and twenty bacterial ASVs were identified as core microbiome members, defined as taxa consistently present across samples and depths over five years **(Table S11)**. Correspondingly, 52 archaeal and 120 bacterial metagenome-assembled genomes (MAGs), corresponding to these ASVs, were identified as core microbiome MAGs, spanning 14 families and accounting for 12.6% of the species-level representative MAGs **(Fig. 7A, Fig. S10 and Table S12;** see **Materials and Methods** for details). Among the bacterial core microbiome, Alphaproteobacteria (n = 70) were predominant, while Bathyarchaeia (n = 19) comprised the majority of the archaeal components **(Fig. 7A and Fig. S10)**. Seven bacterial MAGs were identified as JS1, a typical group of heterotrophic deep-sea microorganisms that mediate carbon cycling in subseafloor environments (*49*) and are particularly abundant in methane-rich marine sediments (*50*). Focusing on methanotrophs, core microbiome included aerobic methane-oxidizing bacteria (MOB) Methylococcales and ANME-3 (family Methanosarcinaceae). Both Methylococcales (0.5%) and Methanosarcinaceae (0.3%) exhibited higher abundances in surface and shallow layers compared to deeper layers (0.003% and 0.002%, respectively) **(Fig. S11 and Table S13)**. Notably, while MOB are conventionally considered restricted to surface niches due to oxygen availability (*51*), our observation of Methylococcales at greater depths than previously reported **(Fig. S11 and Table S13)** (*14*, *30*) suggests that they may contribute to methane consumption below the surface.

**Figure 7.**
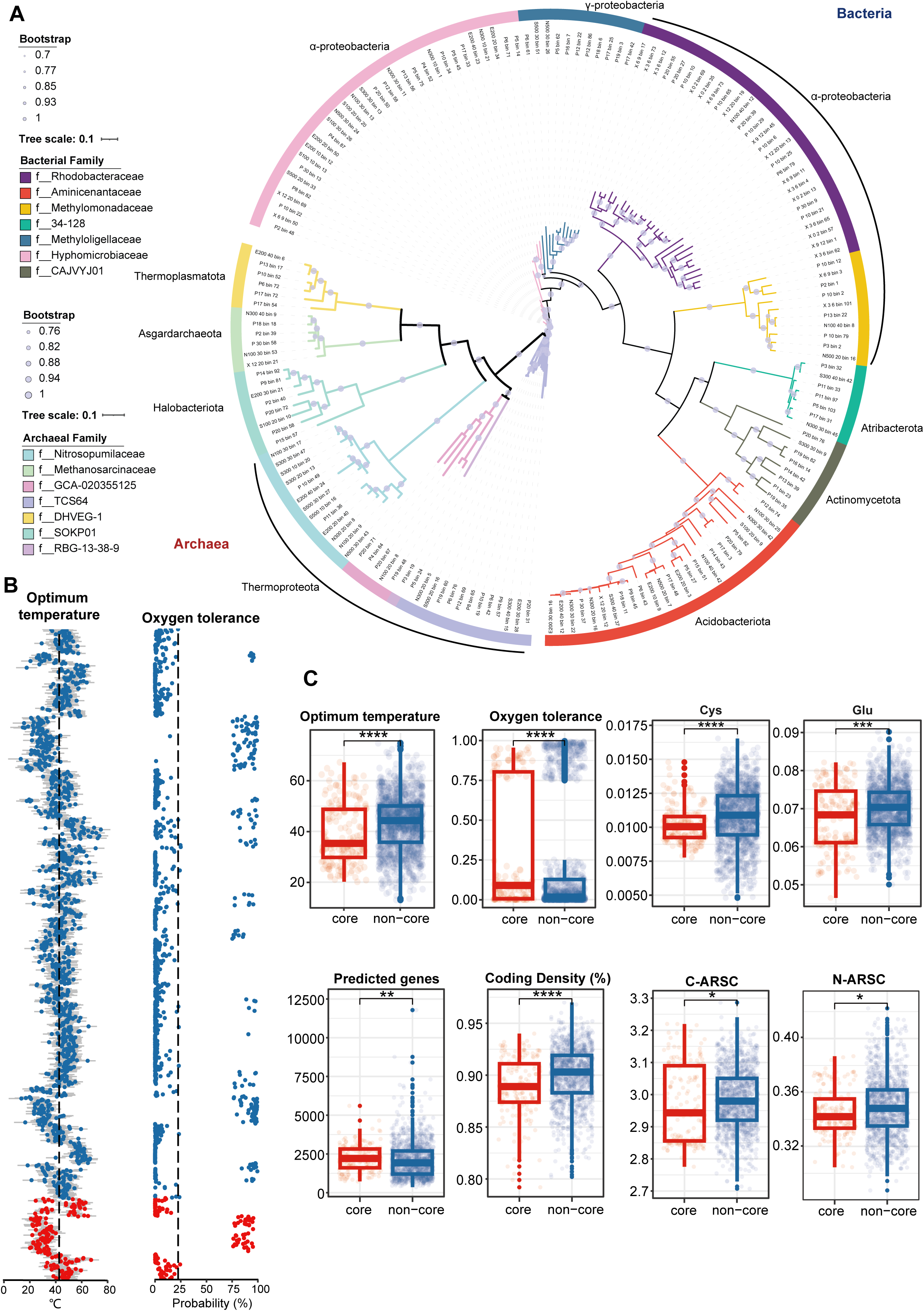
Genomic characteristics of core and non-core microbial communities. **(A)** Maximum-likelihood phylogenomic tree of 172 metagenome-assembled genomes (MAGs) belonging to core microbiome. Source data are provided in **Table S12**. **(B)** Predicted physicochemical growth conditions for each non-redundant MAG, with mean values shown as dots and ±1 the root mean squared error (RSME) shown in lines. The vertical line indicates the average values across all MAGs. For oxygen tolerance, probabilities < 0.5 are classified as ’not tolerant’, while probabilities > 0.5 are classified as ’tolerant’. **(C)** Comparative analysis of predicted growth conditions and genomic traits between core and non-core microbial groups. Genomic traits include predicted gene number, coding density, and C-ARSC and N-ARSC (carbon and nitrogen per amino acid residue side chain). Significance level across groups was statistically checked using Wilcoxon rank-sum tests. Significance levels are denoted as: *, *p* ≤ 0.05; **, *p* ≤ 0.01; ***, *p* ≤ 0.001; ****, *p* ≤ 0.0001. Red represents core microbiome and blue represents non-core microbes. Source data are provided in **Tables S14** and **S15**.

Cold seeps are characterized by low temperatures, typically around 4°C (*2*). The physicochemical growth conditions of microorganisms were predicted based on genomic characteristics, revealing that core microorganisms had significantly lower optimum growth temperatures compared to non-core microorganisms (*p* ≤ 0.0001, **Fig. 7, B and C**). Among core microorganisms, levels of amino acids, such as cysteine (Cys), glutamic acid (Glu) and tyrosine (Try), were significantly lower, while methionine (Met) and valine (Val) levels were significantly higher (**Fig. 7C, Fig. S12 and Table S14**). These amino acid profiles may enhance microbial adaptability to cold environments (*52*). Further genomic analysis revealed that core microbiome exhibited a lower coding density but had more genes and spacers, with longer average spacer lengths compared to non-core microorganisms (*p* < 0.01, **Fig. 7C and Fig. S16**). These traits suggest genomic redundancy, aligning with observations in psychrophilic microorganisms (*53*). Additionally, the majority of core microbiome proteins had an average GRAVY (Grand Average of Hydropathy) score of less than 0 (**Fig. S13 and Table S15**), indicating that they were predominantly hydrophilic. This property may enhance protein flexibility, aiding adaptation to cold environments (*54*). At gene level, the cold adaptation gene *cspA* (*55*), encoding cold shock proteins, was present in 58.6% of MAGs (**Table S16**). Phylogenetic analysis indicated closer relationships among *cspA* sequences within core microorganisms **(Fig. S14)**. Genes involved in translation and post-translational processing are also essential for cold tolerance (*53*), and several of these genes, including *infA* and *infB* (*56*), *gyrA*, and *hns*, were enriched in core microbiome, (Odds Ratio, OR > 1 and False Discovery Rate, FDR < 0.05, **Table S16**).

Core microbiome at QDN-W07 also demonstrated greater tolerance to oxygen and salinity. Predictions of physicochemical growth conditions indicated that core microbiome members exhibited higher oxygen tolerance (*p* ≤ 0.0001, **Fig. 7, B and C**). Regarding salt adaptation, the genes *kamA* and *ablB,* which encode enzymes for synthesizing the osmolyte N(ε)-acetyl-β-L-lysine to combat salt stress (*57*), were found more frequently in the core microorganisms than in non-core ones (**Table S16**), with a notable abundance of *kamA* genes (**Fig. S15**). Microorganisms in surface and shallow sediments showed greater tolerance to both oxygen and salinity, likely due to the influence of core microbiome **(Fig. S16)**. Core microorganisms also had lower average numbers of nitrogen and carbon atoms per amino-acid-residue side chain (N-ARSC and C-ARSC), indicating reduced biosynthetic demands for these elements (*58*) (*p* < 0.05, **Fig. 7C**). Additionally, core microorganisms displayed reduced motility, evidenced by a significantly lower prevalence of the *motA* gene, which encodes a bacterial flagellar motor component (*59*) (OR = 0.59, FDR = 0.03), and the *cheX* gene, which encodes a chemotaxis protein (OR = 0.18, FDR = 0.02, **Table S16**). This reduction in motility suggests increased susceptibility of core microbiome to DL.

### Core microorganisms are pivotal in carbon, nitrogen, and sulfur cycles at QDN-W07

Metabolic functions associated with core microbiome members were identified by quantifying the enrichment of specific genes **(Table S16)** and functions **(Table S17)** in core genomes compared to non-core genomes. Genes related to AeOM, including *pmoABC* and *mmoXYZBCD*, were significantly enriched in core microbiome (FDR ≤ 0.05, **Fig. 8A and Fig. S19)**, highlighting AeOM as a key pathway for methane metabolism at QDN-W07. Additionally, methane oxidation relying on particulate methane monooxygenase (pMMO) was the most enriched function (OR = 10.92, FDR = 0.0001; **Fig. S17**). Core microbiome included 27 potential MOB MAGs, characterized by the presence of key genes, such as *pmoA* for pMMO and *mmoX* for soluble methane monooxygenase (sMMO) (*60*, *61*). Specifically, *pmoA* was identified in six MAGs of Methylomonadaceae, and 24 MAGs contained the gene *mmoX* **(Table S18)**, including members from Aminicenantaceae (*n* = 19), Methylomonadaceae (*n* = 4), and JS1 (*n* = 1). Notably, four Methylomonadaceae MAGs contained complete sMMO operons, with two of these also carrying *pmoA*, suggesting that these methanotrophs might switch between two MMO types based on copper availability (*62*). Other functions related to AeOM, such as oxidation of methanol, formaldehyde and formate (*63*), were also enriched in core microbiome (**Fig. 8A**). One of Methylomonadaceae MAGs (N100_40_bin_8) displayed a complete AeOM pathway.

**Figure 8.**
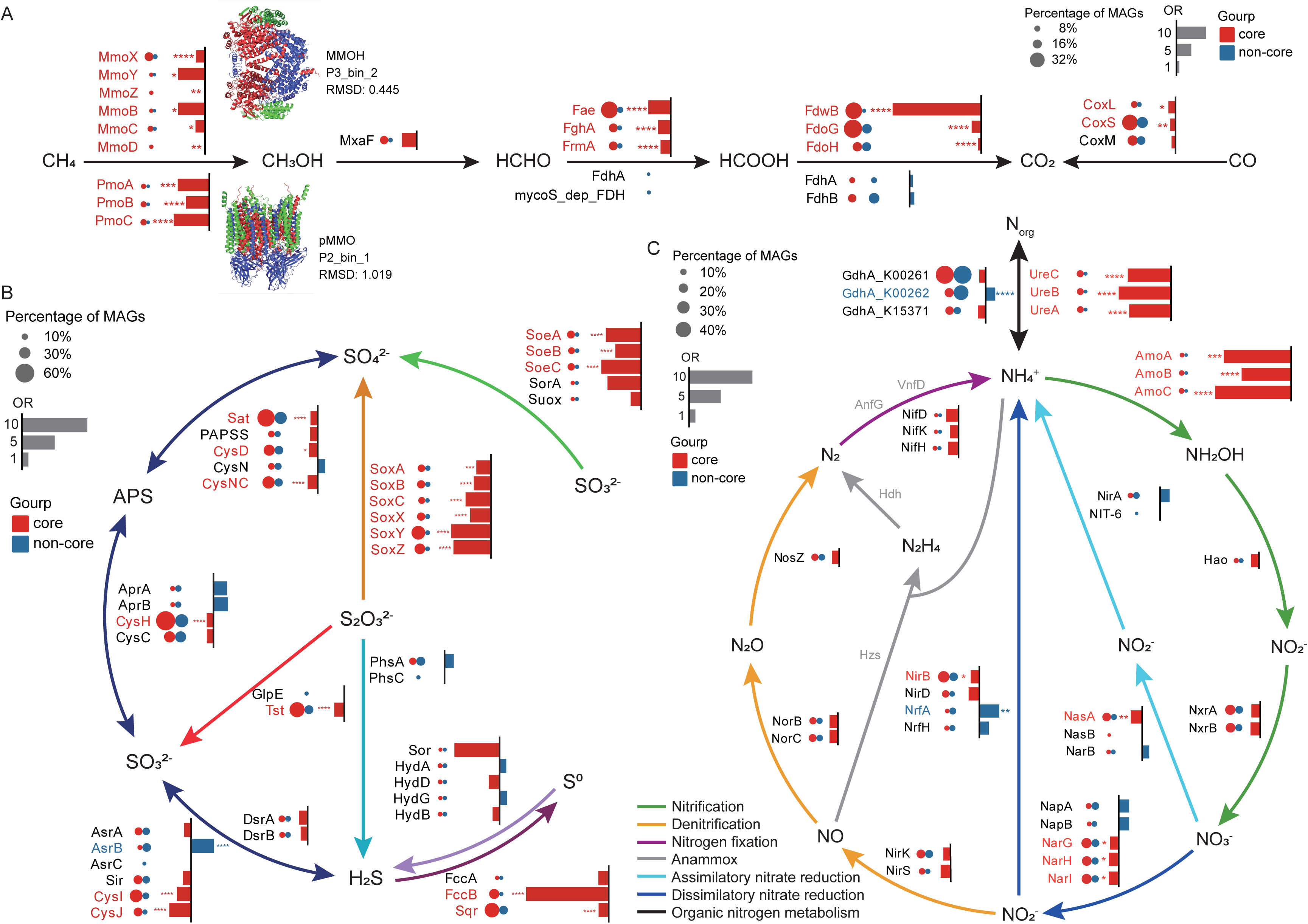
Enrichment of key metabolic genes in core microbiome at QDN-W07. (**A**) Aerobic of methane and CO, (**B**) sulfur and (**C**) nitrogen metabolism. Genes that are more common in the core microbiome are shown in red (OR > 1), while those less common are in blue (OR < 1). Significance levels are denoted as: *, false discovery rate (FDR) ≤ 0.05; **, FDR ≤ 0.01; ***, FDR ≤ 0.001; ****, FDR ≤ 0.0001. In panel (**A**), three-dimensional structures of MMOH and pMMO proteins are shown, with comparisons to reference structures from the PDB database (MMOH, PDB ID: 6CXH; pMMO, PDB ID: 7YZY). Structural similarity is assessed via Root Mean Square Deviation (RMSD) values, where RMSD < 2Å typically indicates a high degree of structural similarity between the two proteins. Source data are provided in **Table S16**.

The gene *mcrA*, essential for AOM (*64*, *65*), was detected in three MAGs from core microbiome and six MAGs from non-core microbiome (**Table S18**). Phylogenetic analysis of *mcrA* **(Fig. S20A)** and taxonomic annotation of these MAGs (**Table S18**) confirmed that all core microbiome *mcrA* originated from ANME-3 (genus *DQIP01*), whereas those in non-core microbiome were associated with ANME-2a/b (family Methanocomedenaceae) (*66*). Predicted structures of methyl coenzyme M reductase (MCR) exhibited high similarity to the reference structure (PDB ID: 1E6V; Root Mean Square Deviation (RMSD) = 0.61∼0.84), further confirming the potential of these microorganisms for AOM (**Fig. S20B**). Overall, core microbiome at QDN-W07 is central to methane metabolism, particularly through AeOM pathways, as evidenced by the enrichment of genes for both pMMO and sMMO. This also highlights the great contributions of MOB and ANME-3 to the microbial community at an early-stage cold seep (*30*).

For sulfur metabolism, the core microbiome exhibited enrichment of genes associated with sulfur oxidation (*fccB* and *sqr*), sulfite oxidation (*soeABC*), and thiosulfate oxidation (*sox*) (**Fig. 8B**). This suggests a substantial presence of sulfur-oxidizing bacteria (SOB) within the core microbiome, which are capable of producing large amounts of sulfate. Such activity could potentially transform sulfate-poor sediments at QDN-W07 **(Fig. 2)** into a sulfate-rich environment. In the nitrogen cycle, genes such as *ure* and *amo* were significantly enriched in core microbiome (FDR ≤ 0.001, **Fig. 8C**). The gene *ureA*, essential for urea metabolism, is widespread in the deep sea and provides a vital nitrogen source for cold seep microorganisms (*67*). Nitrification through ammonia oxidation, facilitated by the enzyme encoded by *amoA*, serves as a primary energy source for carbon fixation in the dark ocean (*68*). At QDN-W07, concentration of NH[[remained noticeably lower than at the other two sites (**Fig. 2 and Table S2**), likely due to extensive utilization by core microorganisms through ammonia oxidation. Additionally, genes associated with aerobic CO oxidation were enriched in core microbiome (**Fig. 8A**), suggesting that CO serves as an important energy source (*69*), providing electron donors for carbon fixation in the deep ocean (*70*). In summary, core microorganisms at QDN-W07 primarily obtain energy through various pathways, including aerobiotic methane oxidation, urea metabolism, ammonia oxidation, and CO oxidation. This metabolic versatility underscores the ecological significance of the core microbiome in maintaining biogeochemical cycles at this cold seep.

## Materials and Methods

### Site description and sampling

Sediment samples were collected from three cold seep sites, QDN-W07, QDN-S18, and HM-ROV01, located within the Haima cold seep situated in the QDNB on the northern slope of the South China Sea **(Fig. 1 and Table S1)**. Specifically, samples from QDN-W07 were collected over five consecutive years from 2019 to 2023.

Sediment samples were collected primarily in three ways: (1) 142 samples from short push cores collected by “Haima” ROV dives during the R/V Haiyangdizhi6 cruise in April 2020, April 2021, November 2021, July 2022, and April 2023; (2) 85 samples from long piston cores collected by gravity piston sampler during the Haiyangdizhi10 cruise in June 2019, the Haiyangdizhi4 cruises in March 2022 and June 2023; (3) five samples from box cores during the Haiyangdizhi6 cruise in November 2021. Samples collected by push core and box core were generally collected at depths from 0 to 60 cmbsf, representing surface and shallow layers. In contrast, the piston cores enabled collection of deep layer samples, reaching up to 700 cmbsf. Upon retrieval, the piston core and push core were immediately cut into segments on board at the interval of 20 and 10 cm from top to bottom, respectively. The box core was separated by three centimeters. Pore water was extracted from sediments by Rhizon samplers (0.2 μm pore size; Rhizosphere) as soon as the core arrived on deck for geochemical analyses. All sediment samples were stored at -80 °C until DNA extraction. Based on depth, sediment samples were classified into three distinct groups (**Table S1**): surface (n = 44), shallow (n = 121), and deep (n = 67). Surface samples were collected from the topmost sediment layers, shallow samples from depths under 80 cmbsf, and deep samples from depths exceeding 80 cmbsf.

### High-resolution 3D seismic survey

A high-resolution 3D seismic survey was utilized to analyze the subsurface structures of the newly discovered cold seep, QDN-W07, in the central QDNB. The 3D seismic data were acquired in 2015 by the China National Offshore Oil Corporation and processed as previous study (*20*). It was acquired using 12 parallel streamers with 100 m spacings. The data have inline (NW-SE) and crossline (SW-NE) spacings of 12.5 m and 12.5 m, respectively and 1.0 ms sampling interval.

### Physical and geochemical analyses

We measured a total of 17 environmental factors using 235 pore water samples (**Table S2**). Total alkalinity (TA), PO_4_^3-^, NH_4_^+^, Mn^2+^, Fe^2+^, CH_4_, δ^13^C_DIC_, H_2_S, SO_4_^2-^, calcium ion (Ca^2+^), and magnesium ion (Mg^2+^) were determined as previously described (*27*, *71*). The pH of the pore water was monitored using a calibrated pH meter. NO_2_^-^ was determined using colorimetric techniques like the Griess assay. In this method, pore water samples were mixed with sulfanilic acid and *N*-(1-naphthyl)ethylenediamine dihydrochloride, and the resulting azo dye was measured spectrophotometrically at 543 nm. NO_3_^-^ was first reduced to NO_2_^-^ using cadmium-coated zinc strips, and the reduced NO_2_^-^ was then detected using the Griess reaction as described above. Concentration and isotopic analyses of DIC were analyzed using a multiflow-isotope ratio mass spectrometer (Thermofisher, Delta V Advantage, USA), with an analytical precision of 0.2‰. Concentrations of chloride (Cl^-^) and bromide (Br^-^) were measured by an ICS-1100 ion chromatography (Thermofisher, USA) with an analytical error of ± 1%. Concentrations of Na^+^ and K^+^ were determined with the same ion chromatography system, ensuring an analytical precision of less than 10%.

### DNA extraction, 16S rRNA amplicon sequencing and metagenomic sequencing

DNA extractions were performed for 16S rRNA amplicon sequencing from 232 sediment samples, and metagenomic sequencing from 54 samples **(Table S1)**. DNA was extracted from sediment samples using the PowerSoil DNA Isolation Kit (MoBio Laboratories, Carlsbad, CA, USA) following the manufacturer’s instructions. We used 0.5 g of each sediment sample for 16S rRNA amplicon sequencing and 10 g for metagenomic sequencing. Amplicon sequencing was performed via the Illumina Miseq PE300 (Illumina, Inc., San Diego, CA, USA) using a paired-end approach. Specific primer pairs were used to target different domains of life. The primer pair 338F (5′-ACTCCTACGGGAGGCAGCAG-3′) and 806R (5′-GGACTACHVGGGTWTCTAAT-3’) was used to amplify the V3-V4 region of the bacterial 16S rRNA gene. The V3-V5 regions of the archaeal 16S rRNA gene were amplified using the primer pair Arch344F (5′-ACGGGGYGCAGCAGGCGCGA-3′) and Arch915R (5′-GTGCTCCCCCGCCAATTCCT-3′). Metagenomic sequencing was performed via the Illumina NovaSeq 6000 (Illumina, Inc., San Diego, CA, USA) using a paired-end approach.

### 16S rRNA amplicon sequencing data analysis

The sequencing data were then processed following the DADA2 pipeline (*72*) within QIIME 2 platform (*73*), which includes filtering, denoising, paired-end sequence merging, chimera filtering, and generation of ASVs. Sequence reads were rarefied to equal depths per sample, with 12,328 reads for bacteria and 6,473 reads for archaea. Taxonomic assignment was performed using the SILVA v138.1 database (*74*), with a confidence threshold of > 0.7. Shannon index was calculated with mothur (v1.30) (*75*) to evaluate alpha-diversity of microbial communities.

Microbial network analysis was performed to explore the co-occurrence patterns and potential interactions among microbial taxa. Based on relative abundance of archaea and bacteria ASVs, co-occurrence networks of prokaryotic communities within the seepage areas across the three sites were constructed using the Integrated Network Analysis Pipeline (iNAP, http://mem.rcees.ac.cn:8081) (*76*) on the Galaxy platform (v21.01) (*77*). Samples were grouped by site and year into six distinct groups, retaining only ASVs present in all samples within each group. We used FastSpar (*78*) to implement the SparCC (*79*) algorithm to calculate correlation values with default parameters. Only significant correlations (|*R*| ≥ 0.6 and *p* < 0.05) were retained and visualized in Gephi (v0.10.1) (*80*). Network characteristics, such as average degree, network density and network diameter, were also calculated in Gephi.

### Community assembly processes

The contribution of different ecological processes to community assembly was quantified using the phylogenetic-bin-based null model implemented with R package iCAMP (*38*). Within the framework, community assembly processes were attributed to deterministic assembly, including HoS and heterogeneous selection (HeS), and stochastic assembly, including homogenizing dispersal (HD), DL and ‘drift’ (DR). In this context, ‘drift’ represents drift, diversification, weak selection, or weak dispersal.

First, the observed taxa were divided into different phylogenetic groups (bins) according to a phylogenetic signal threshold (*ds* = 0.2) within the phylogenetic tree constructed using FastTree (*81*) (parameters: -nt -gtr -gamma). If a bin contained fewer ASVs than the minimum size requirement (*n_min_*), smaller bins were merged with the nearest relatives until the *n_min_* threshold was met. Using environmental variables, the dniche and ps.bin functions in the iCAMP package were employed to calculate within-bin phylogenetic signals to determine *n_min_* value. After testing values from 24 to 72, we selected *n_min_* values that maximized RAsig.abj (relative abundance of bins with significant phylogenetic signals) and yielded relatively high meanR (mean correlation coefficient across bins). Ultimately, *n_min_* was set to 42 for archaea and 30 for bacteria. Subsequently, based on the null-model analysis, the beta-net relatedness index (βNRI) and modified Raup-Crick metric (RC) were calculated for all the bins between each pair of samples to identify the assembly processes governing each bin’s turnovers. Mantel tests with constrained permutations were performed to assess the effects of environmental factors on HoS. Finally, the fraction of individual processes across all bins were weighted by the relative abundance of each bin and summarized to estimate the relative importance of individual processes at the whole community level. The icamp.cate function was used to assess the relative importance of each process to the core microbiome and the subcommunity at different taxonomic levels.

### Metagenomic sequence data analysis

Metagenomic sequence data were quality controlled using fastp (*82*) (v0.23.2; default parameters). Clean reads from each cold seep sediment metagenome were then assembled using MEGAHIT (*83*) (v1.2.9; default parameters). The binning process was performed with the metaWRAP (*84*) binning module (v1.3.2; parameters: - metabat2, -maxbin2, -concoct, -universal). The bins obtained from each binning tool were subsequently integrated and refined using the Bin_refinement module of the metaWRAP pipeline (v1.3.2; parameters: -c 50 -x 10). The completeness and contamination of refined bins were evaluated with CheckM (*85*) (v1.2.1). All MAGs were dereplicated at the species level using dRep (*86*) (v3.4.0; parameters: -comp 50 -con 10) with an average nucleotide identity (ANI) cutoff value of 95%. Representative genomes were selected based on dRep scores, which were derived from genome completeness and contamination, resulting in a final set of 1365 non-redundant MAGs. MAGs were taxonomically classified using the GTDB-Tk (v2.3.2) (*87*) with default parameters against the GTDB R214 database (*88*). The coverage of each MAG was calculated using CoverM in genome mode (v0.6.1; https://github.com/wwood/CoverM; parameters: -min-read-percent-identity 0.95 - min-read-aligned-percent 0.75 -trim-min 0.10 -trim-max 0.90 -m relative_abundance) by mapping clean reads from the 54 metagenomes to all MAGs.

16S rRNA gene sequences were extracted from MAGs with Phyloflash v3.4 (*89*) and Barrnap (https://github.com/tseemann/barrnap). Specifically, Phyloflash was used to assemble and reconstruct clean reads into full-length 16S rRNA gene sequences. We then used markerMAG (*90*) to associate the full-length 16S rRNA gene sequences with their corresponding MAGs, retaining only the linked sequences (with identity > 99%) for subsequent analyses. This process resulted in a total of 599 bins matched to the assembled full-length 16S rRNA sequences. Barrnap was used to predict the location of 16S rRNA genes within MAGs. 16S rRNA sequences were identified from 331 MAGs, of which 118 were full-length 16S rRNA gene sequences.

### Criteria for core microbiome at QDN-W07

Samples from the seepage area were divided into eight groups based on year and depth, aiming to identify prokaryotes that were persistently present across different spatiotemporal conditions at QDN-W07. Specifically, samples collected from 2019 to 2023 were separated into two depth ranges each year: < 60 cmbsf and > 60 cmbsf. However, for 2020 and 2021, only samples from the < 60 cmbsf range were obtained, resulting in a total of eight groups. The core microbiome was operationally defined as microbial taxa detected across all eight sample groups, based on the ASV table.

Two methods were used to identify the MAGs representing the core microbiome. The first method involved matching 16S rRNA gene sequences obtained from metagenome and amplicon sequencing. Specifically, 16S rRNA gene sequences extracted from MAGs using Phyloflash and Barrnap were aligned against the core microbiome 16S rRNA amplicon sequences using BLASTN. Only alignments with an identity > 99% and an alignment length > 270 bp for archaea and > 400 bp for bacteria were retained. The second method involved identifying MAGs taxonomically congruent with the core microbiome at the family level. Due to naming discrepancies between the Silva database and the GTDB R214 database, 16S rRNA gene amplicons representing the core microbiome were reclassified using the GTDB SSU and GreenGenes2 databases (*91*). In detail, target sequences were initially extracted from the reference databases based on matches to the primer pairs used for amplicon sequencing. Then, two Naive Bayes classifiers were trained on the extracted reads and taxonomy from each reference database. Finally, the 16S rRNA gene amplicons were classified using these trained classifiers, with annotations filtered at a 99% confidence threshold. The feature-classifier tool within QIIME 2 (v2022.8) (*73*) was used to process this classification workflow.

### Genomic characteristics and metabolic function

GC content, gene number, genome size and coding density were calculated using CheckM (*85*). Specifically, predicted genome size was determined using the formula: original assembled MAG size / (completeness + contamination) (*92*). C-ARSC, N-ARSC, and intergenic spacer lengths were calculated by the python scripts “get_ARSC.py” and “calculate_intergenic_spacers.py” (https://github.com/faylward/pangenomics/), respectively. The hydrophobicity and aromaticity indices of predicted gene sequences for each MAG were calculated using CodonW (v1.4.4, https://codonw.sourceforge.net/), with averages taken to represent the overall hydrophobicity and aromaticity of the MAGs. Amino acid compositions of MAGs were calculated by the predicted proteins of each MAGs. GenomeSPOT (*93*) was used to predict microbial growth conditions, including salinity, pH, oxygen tolerance, and temperature based on genomes.

Metabolic pathway prediction was conducted using the METABOLIC pipeline (v4.0) (*94*) and eggNOG-mapper (v2) (*95*, *96*). Phylogenetic trees were constructed for key genes and 16S rRNA gene amplicon sequences of ANME. The analyzed key genes included methane oxidation genes (*pmoA*, *mmoX*, and *mcrA*), a cold shock protein gene (*cspA*), and salt adaptation genes (*kamA* and *ablB*). Reference sequences for *pmoA* and *mmoX* were obtained from the Greening lab metabolic marker gene databases (*97*), while *mcrA* sequences were sourced from Woods’ study (*66*) and other references from Swiss-Prot (*98*). Sequences were aligned using MUSCLE (v3.8.1551) (*99*) and trimmed with TrimAL (v1.4.1; parameters: -gappyout) (*100*). Maximum-likelihood trees were constructed with IQ-TREE (v2.2.0.3; parameters: -m MFP -bb 1000) (*101*). The trees were visualized and refined using Interactive Tree of Life (iTOL; v6) (*102*). Alphafold 3 (*103*) was applied to predict the structure for MCR, pMMO and MMOX. Reference protein structures of these proteins were downloaded from the Protein Data Bank (PDB) (*104*). All structures were visualized and exported as images using PyMOL (*105*).

### Statistical analysis

Statistical analyses were conducted using R (v4.2.3). Principal coordinates analysis (PCoA) was performed based on weighted Unifrac distance. Depth decay relationship (DDR) was calculated as slopes of ordinary least-squares regressions for the relationships between depths or environment factors and community similarities (1 - dissimilarity of the Bray-Curtis metric). After dropping missing values, 13 environmental factors were used in statistical analyses. Mantel correlations were calculated and visualized using R package linkET (*106*). Distance-based redundancy analysis (db-RDA) was performed based on Bray-Curtis distance with R package vegan (https://vegandevs.github.io/vegan/) and usdm. A total of 999 permutations were performed to evaluate the significance. Chi-squared tests and Fisher tests evaluated the over/under enrichment of each identified function category in core microbiome genomes relative to all MAGs using the chisq.test (expected > 5) and fisher.test (expected ≤ 5) function in R. *P*-values were adjusted for multiple testing using false discovery rate with the p.adjust function in R, with an FDR ≤ 0.05 considered statistically significant. Odds ratio (OR) was calculated to assess the strength of the relationship between genes or function and core microbiome.

## Acknowledgments

We thank Qiuyun Jiang, Chuwen Zhang, Jiaxue Peng, Yongyi Peng, Minghui Geng and Xuemin Wu for providing valuable comments.

## Author contributions

Conceptualization: XD

Methodology: XD, XL, XX

Investigation: XL, XX, QL, JZ, YD, BG

Visualization: XL, XX, TC, JW, YH

Supervision: XD, QL, XX

Writing—original draft: XL, XX, XD, QL

Writing—review & editing: XD, XL, XX, QL, LZ, JW, SL

## Data and materials availability

All sequences of 16S rRNA gene amplicon and metagenome obtained in this study have been deposited in the National Center for Biotechnology Information (NCBI) under BioProject accession number PRJNA1169195.

