## Supplemental Figures 1-20 for "Microbial succession, community assembly and adaptation over five years in a newly discovered deep-sea cold seep"

This PDF file includes:

Figs. S1 to S20

Legends for tables S1 to S18

Other Supplementary Materials for this manuscript include the following:

Tables S1 to S18

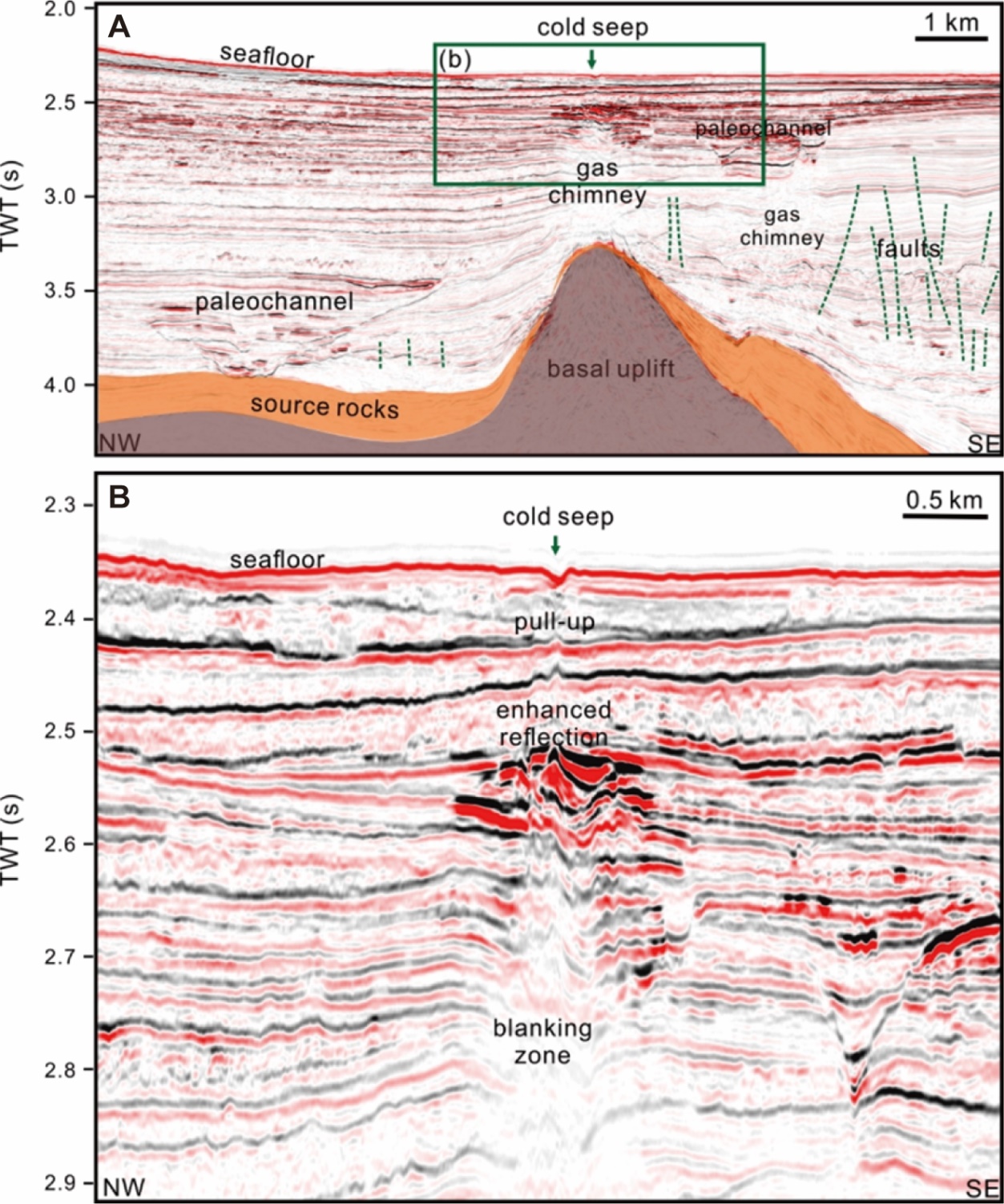

Figure S1. Seismic reflection profiles showing the geological features associated with the newly discovered cold seep in the study area. (A) A seismic profile illustrating large-scale geological features from the seafloor to the basal uplift. (B) A zoomed-in view of the seismic profile (green box in panel (A)) showing detailed features near QDN-W07.

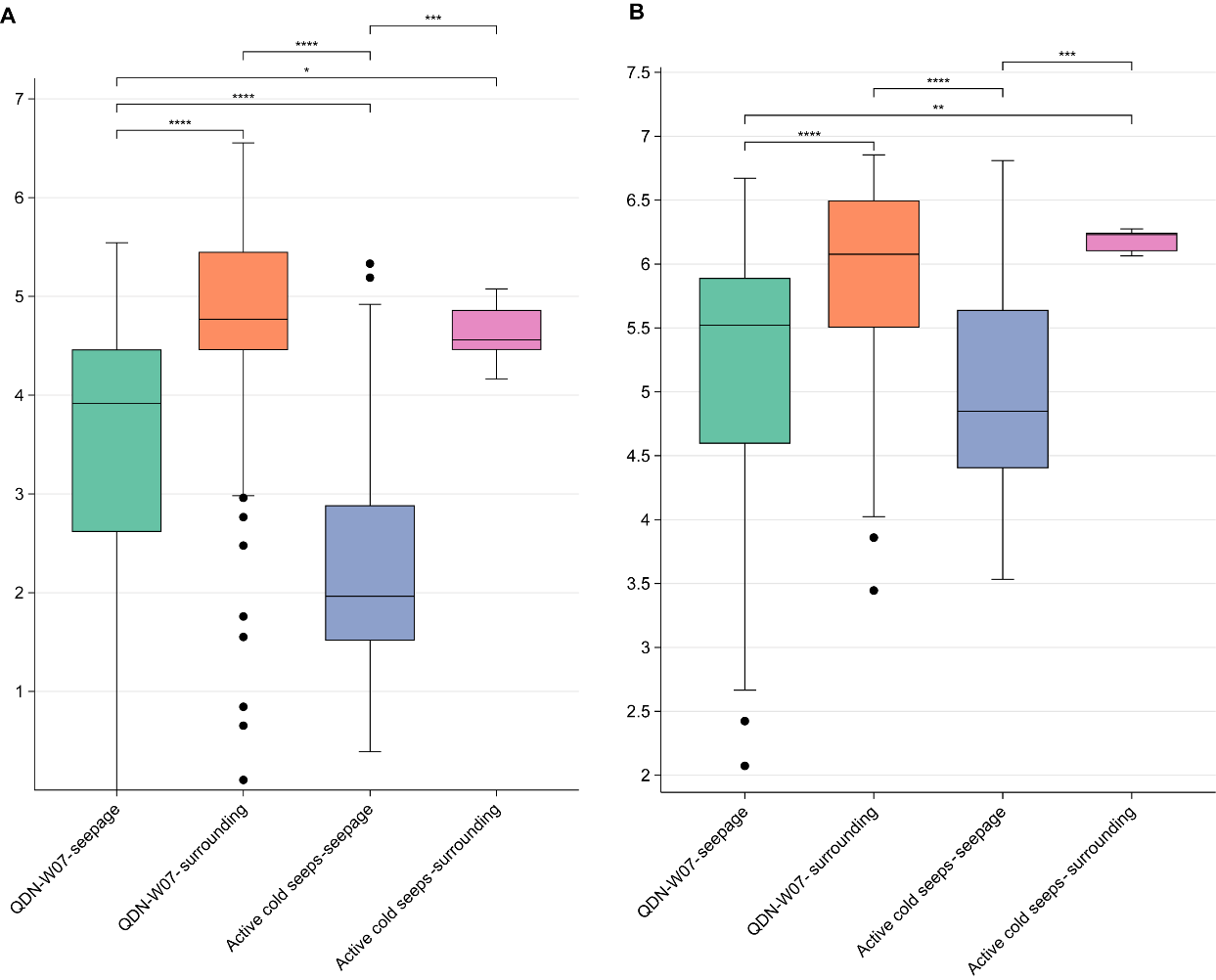

Figure S2. Alpha diversity of microbial communities at different cold seep sites. Alpha diversity is represented by Shannon indices of (A) archaeal and (B) bacterial communities. Samples are grouped by sites and distance from the seep (seepage or surrounding area). Active cold seeps include HM-ROV01 and QDN-S18. Statistical significance of median differences across groups was assessed using Wilcoxon rank-sum tests. Significance levels are denoted as follows: *****p* < 0.0001, ****p* < 0.001, ***p* < 0.01, **p* < 0.05.

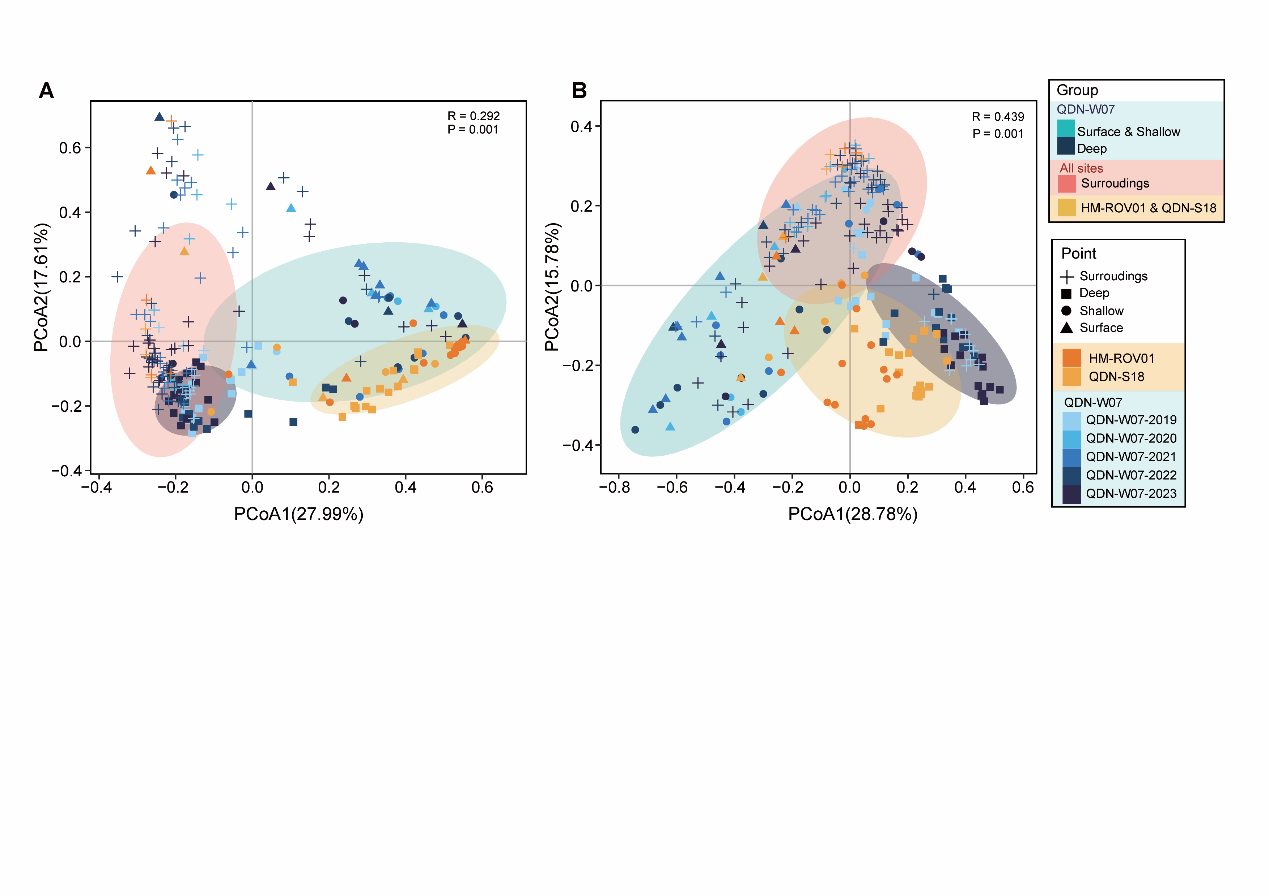

Figure S3. Microbial community structure in cold seep sediments. (A-B) Principal coordinates analysis (PCoA) of archaeal (A) and bacterial (B) communities, colored by sample groups. Differences between groups were analyzed using ANOSIM.

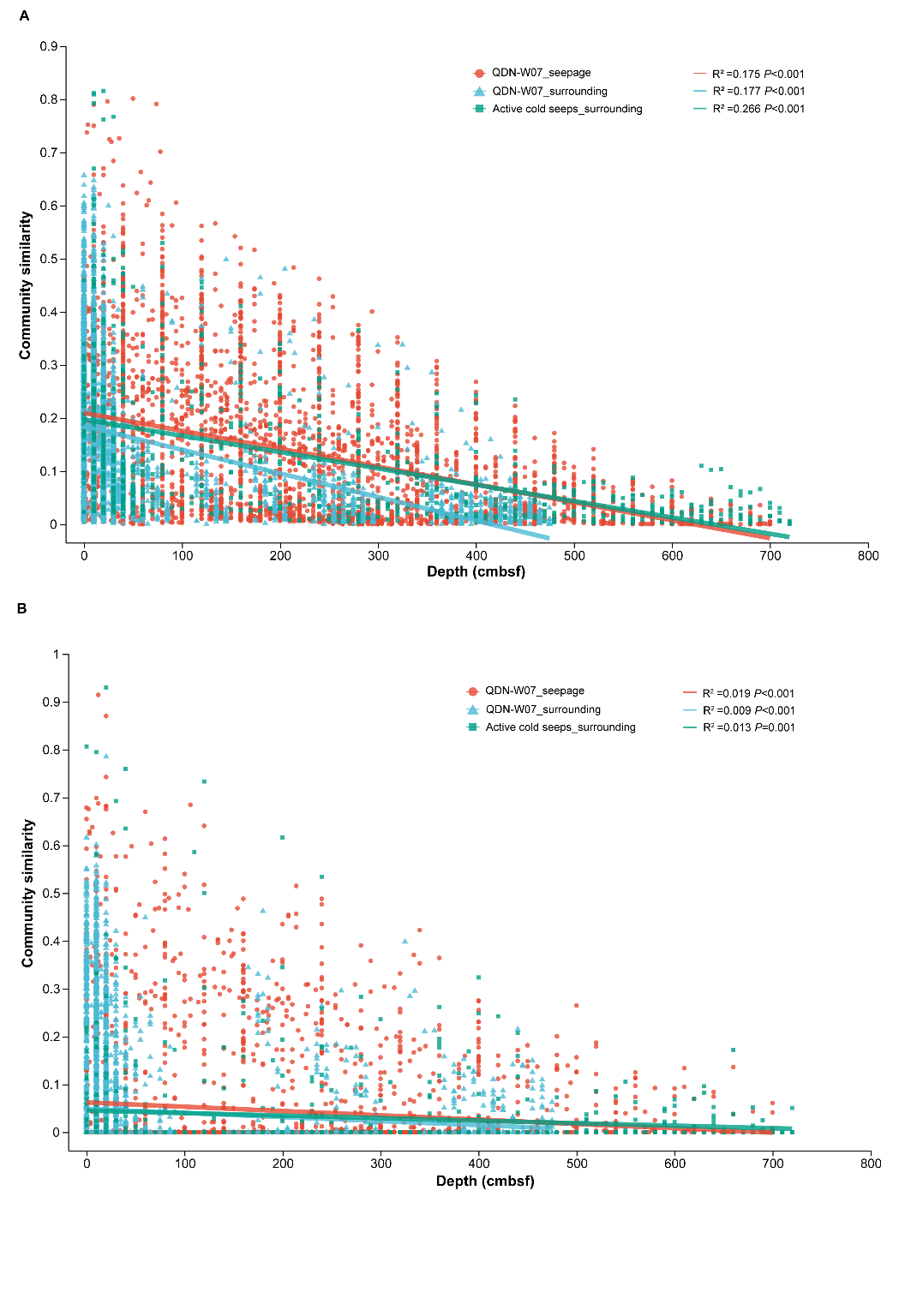

Figure S4. Depth-decay pattern across different cold seep sites. Pairwise relationships between community similarity (1 - Bray-Curtis dissimilarity) and sediment depth are shown for (A) bacteria and (B) archaea. Active cold seeps include HM-ROV01 and QDN-S18. Spearman’s rank correlations were used to assess these relationships.

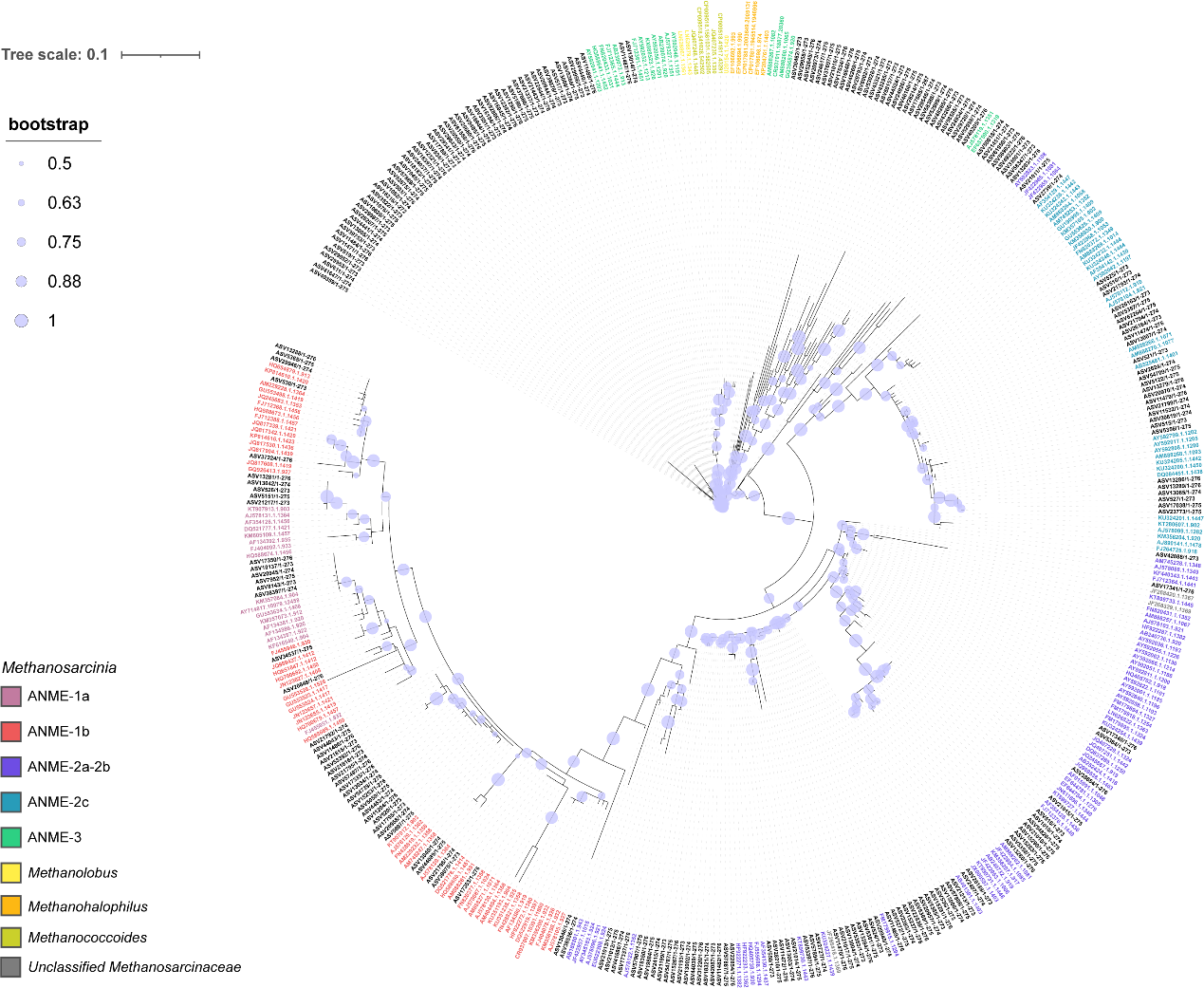

Figure S5. Phylogenetic tree of Methanosarcinia based on 16S rRNA gene sequences. Nodes with colored labels represent reference sequences collected from SILVA, while nodes with black labels represent the amplicon sequence variants (ASVs) identified in this study. Phylogenetic relationships were inferred to highlight the placement of these ASVs within known ANME lineages. Bootstrap values over 50% were shown next to the nodes.

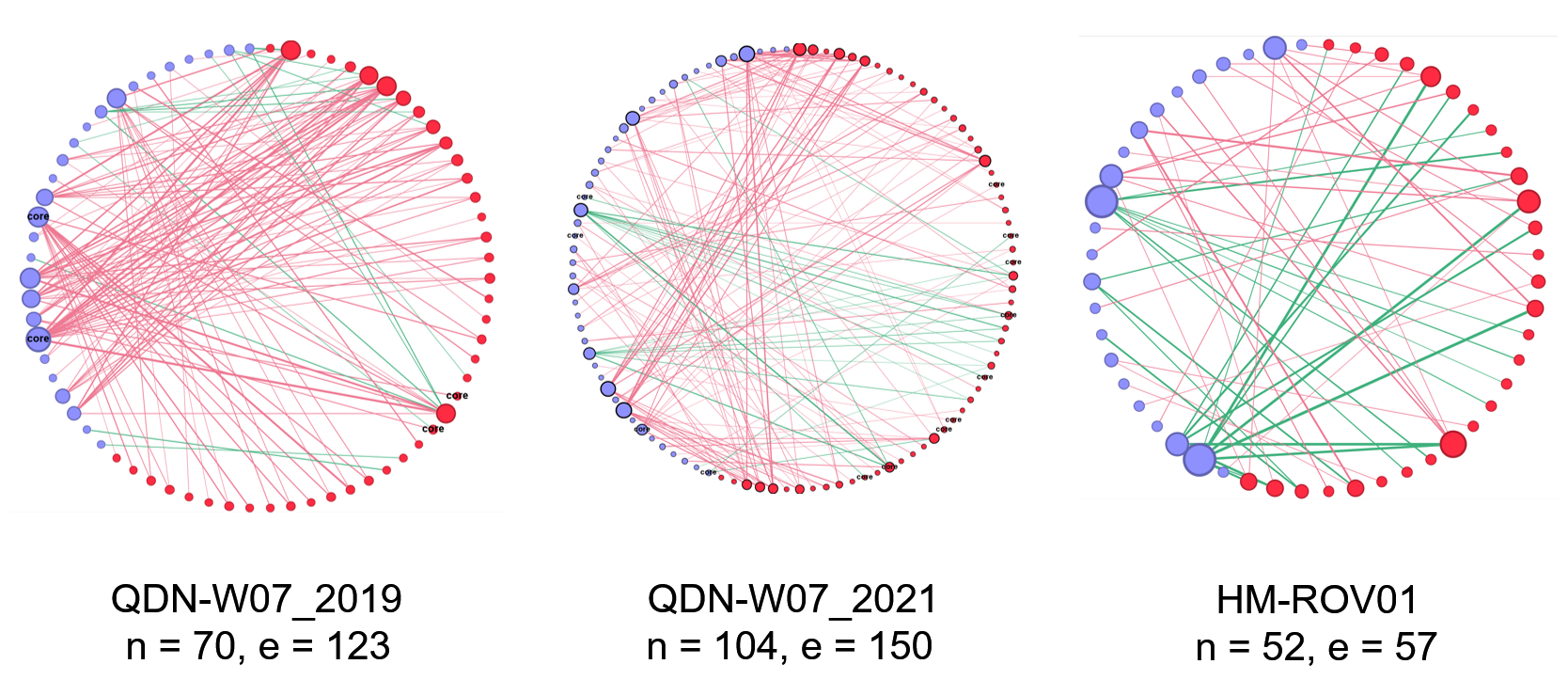

Figure S6. Inter-domain ecological networks (IDENs) for archaea and bacteria at QDN-W07 (in 2019, 2021) and HM-ROV01. In these networks, nodes represent microbial species and edges indicate interactions. Nodes are colored by domain, with red nodes representing archaea and blue nodes representing bacteria. The size of nodes shows their degree of connectivity. Red edges represent significant positive correlations and green edges represent significant negative correlations. Parameters include: n: the number of nodes; e: the number of edges.

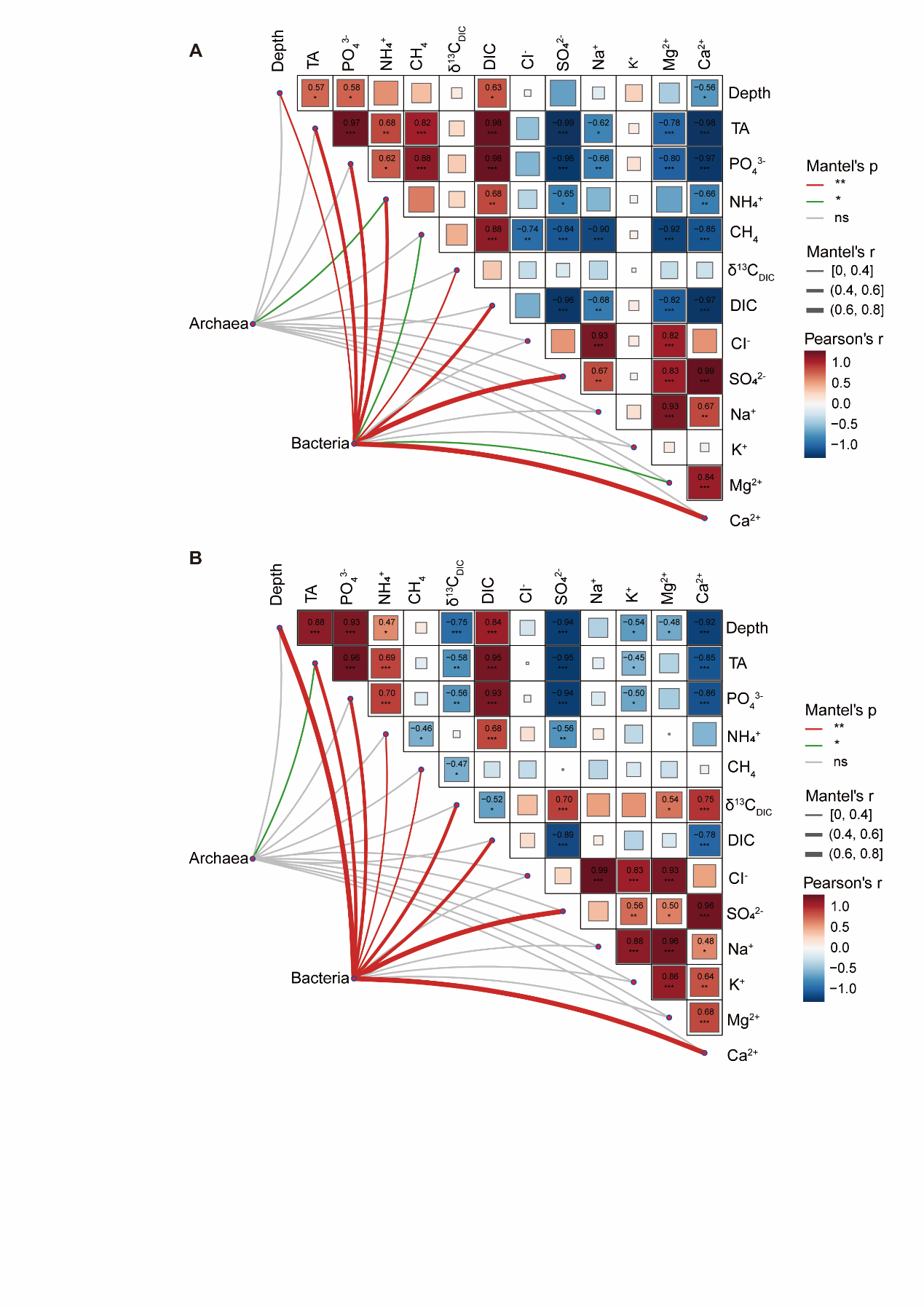

Figure S7. Mantel's correlation analysis between geochemical data and microbial communities at two active cold seep sites. (A) Analysis at HM-ROV01 and (B) analysis at QDN-S18. Edge width corresponds to Mantel’s r statistic and edge color denotes statistical significance. Pairwise comparisons of geochemical data are visualized with a color gradient representing Pearson’s correlation. Source data are provided in Table S6.

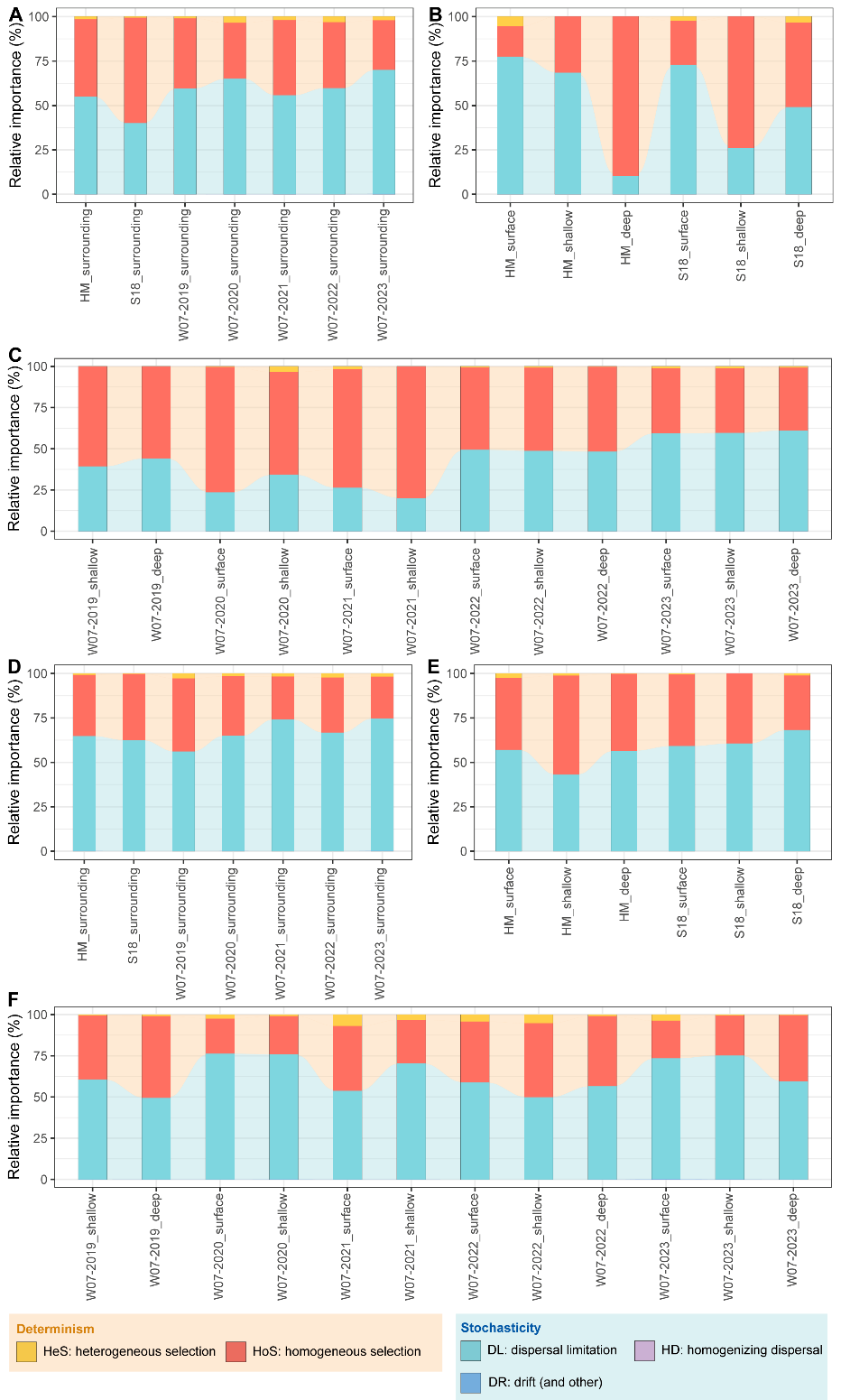

Figure S8. Microbial community assembly processes at QDN-W07 (W07), QDN-S18 (S18) and HM-ROV01 (HM). Relative importance of five ecological processes for (A-C) archaeal and (D-F) bacterial communities across different samples. (A) and (D) show community assembly processes in samples from surrounding areas of cold seeps. (B) and (E) show community assembly processes at varying sediment depths at S18 and HM. (C) and (F) show community assembly processes in W07 seepage area samples collected from different sediment depths from 2019 to 2023. Source data are provided in Table S7.

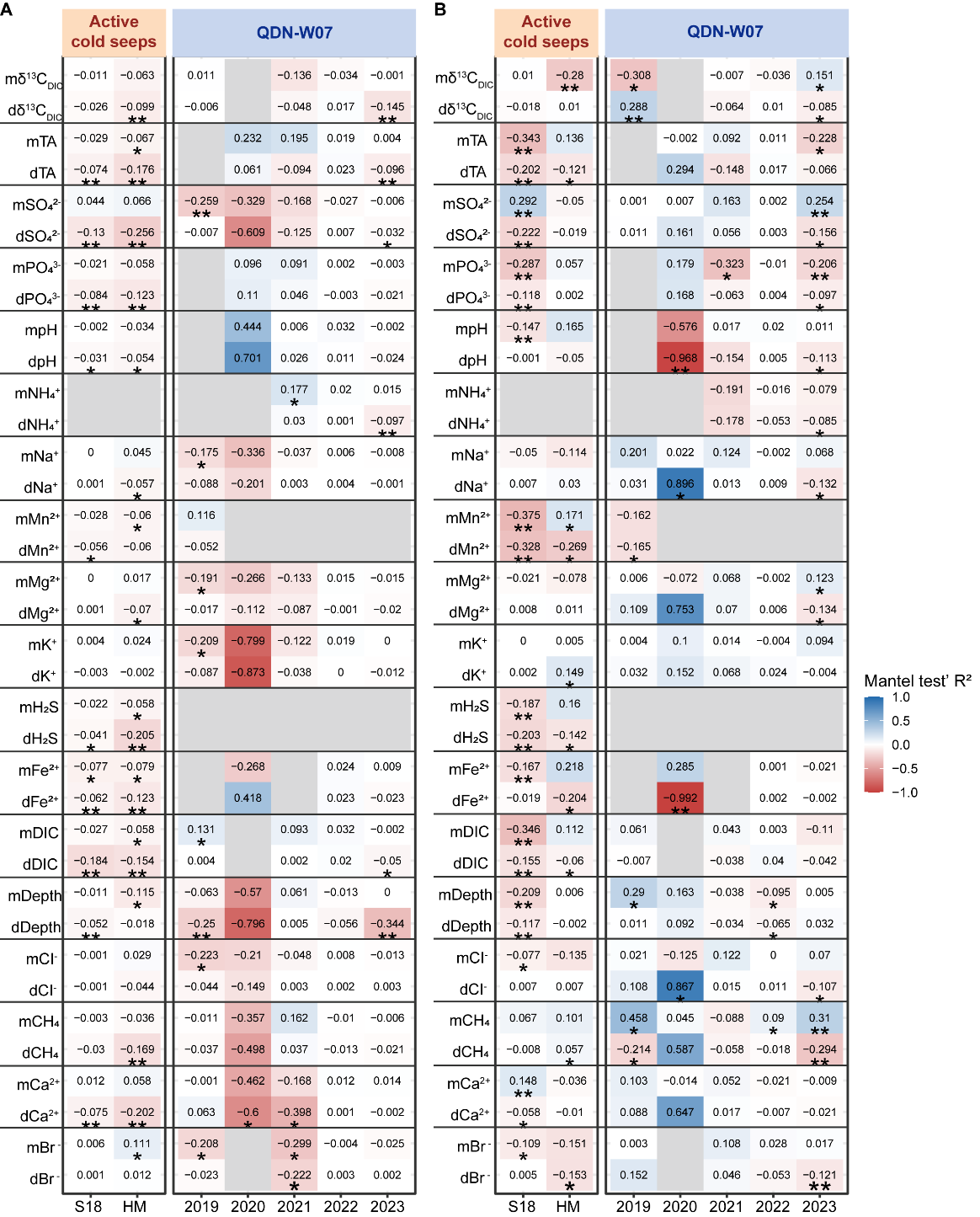

Figure S9. Effects of environmental factors on homogeneous selection (HoS). (A-B) Correlations between environmental factors and HoS for (A) archaea and (B) bacteria, based on Mantel tests. Correlations were determined based on the difference (with a “d” before factors) or the mean (with a “m” before the factors) of a factor between each pair of samples. Colored squares represent the strength and direction of each correlation, with asterisks denoting statistical significance (*, *p* ≤ 0.05; **, *p* ≤ 0.01; ***, *p* ≤ 0.001). Source data are provided in Table S8.

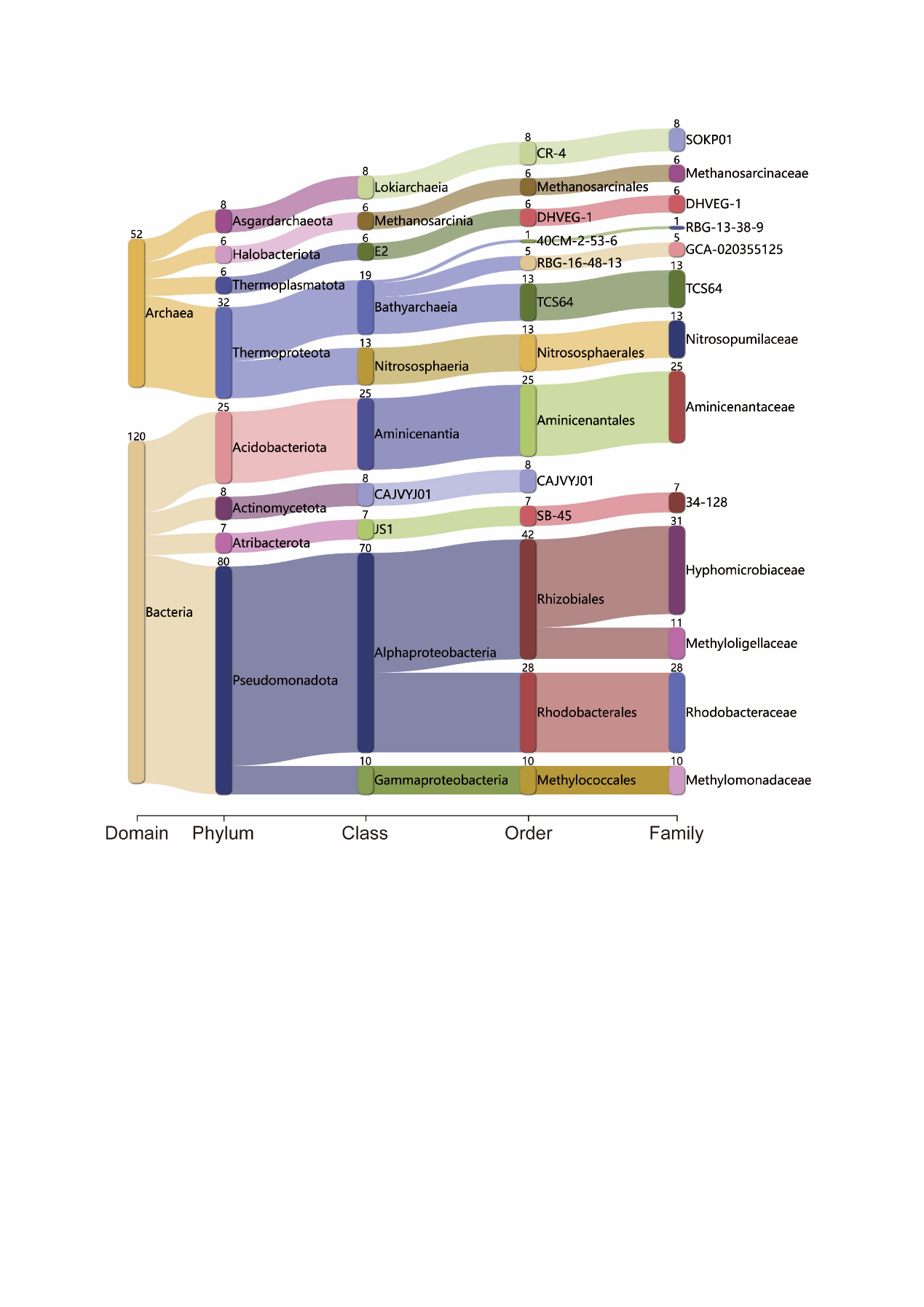

Figure S10. Composition of core cold seep microbiome at QDN-W07. The sankey plot shows MAGs of QDN-W07 core microbiome at different taxonomic levels based on the GTDB-Tk classification. The numbers indicate the number of MAGs recovered for the lineage. Source data are provided in Table S12.

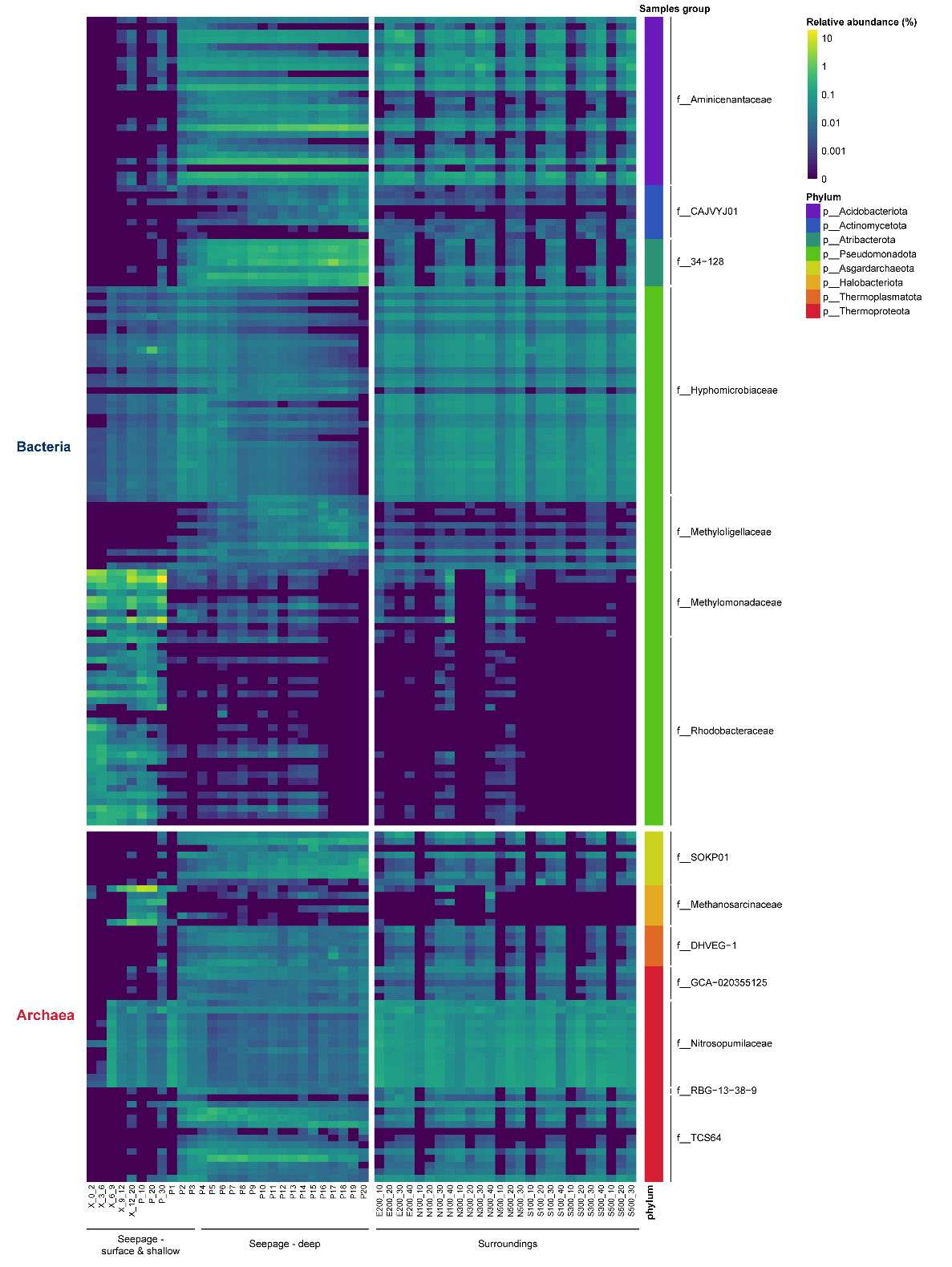

Figure S11. Relative abundance of core microbiome at QDN-W07. Samples were grouped based on their depth and location within the seepage area or surrounding area. Source data are provided in Table S13.

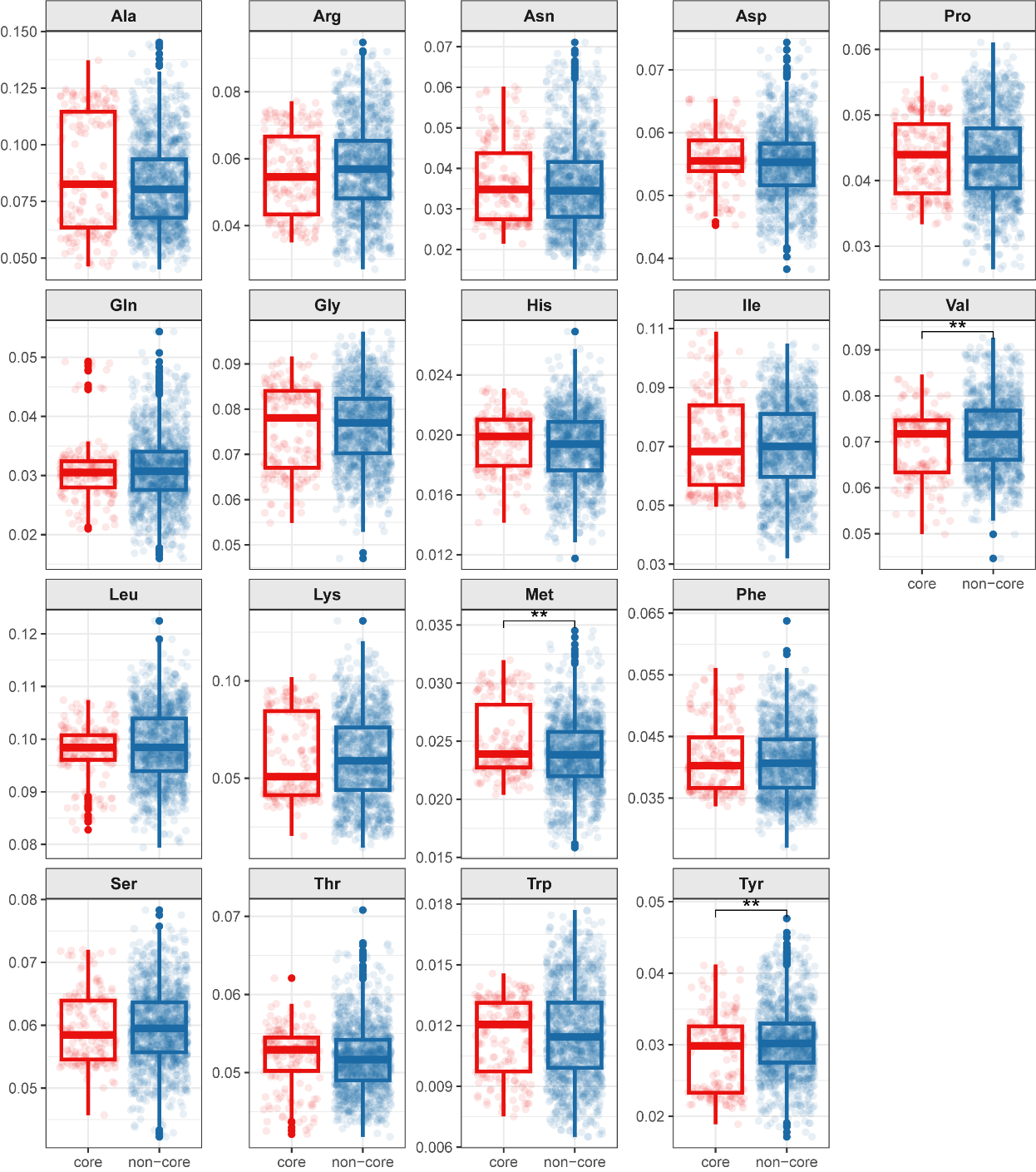

Figure S12. Comparison of amino acid compositions between MAGs from core and non-core microorganisms at QDN-W07. The significance level across groups was assessed using Wilcoxon rank-sum tests. Significance levels are denoted as: *, *p* ≤ 0.05; **, *p* ≤ 0.01; ***, *p* ≤ 0.001; ****, *p* ≤ 0.0001. Ala, Alanine; Arg, Arginine; Asn, Asparagine; Asp, Aspartic acid; Pro, Proline; Gln, Glutamine; Gly, Glycine; His, Histidine; Ile, Isoleucine; Val, Valine; Leu, Leucine; Lys, Lysine; Met, Methionine; Phe, Phenylalanine; Ser, Serine; Thr, Threonine; Trp, Tryptophan; Tyr, Tyrosine. Source data are provided in Table S14.

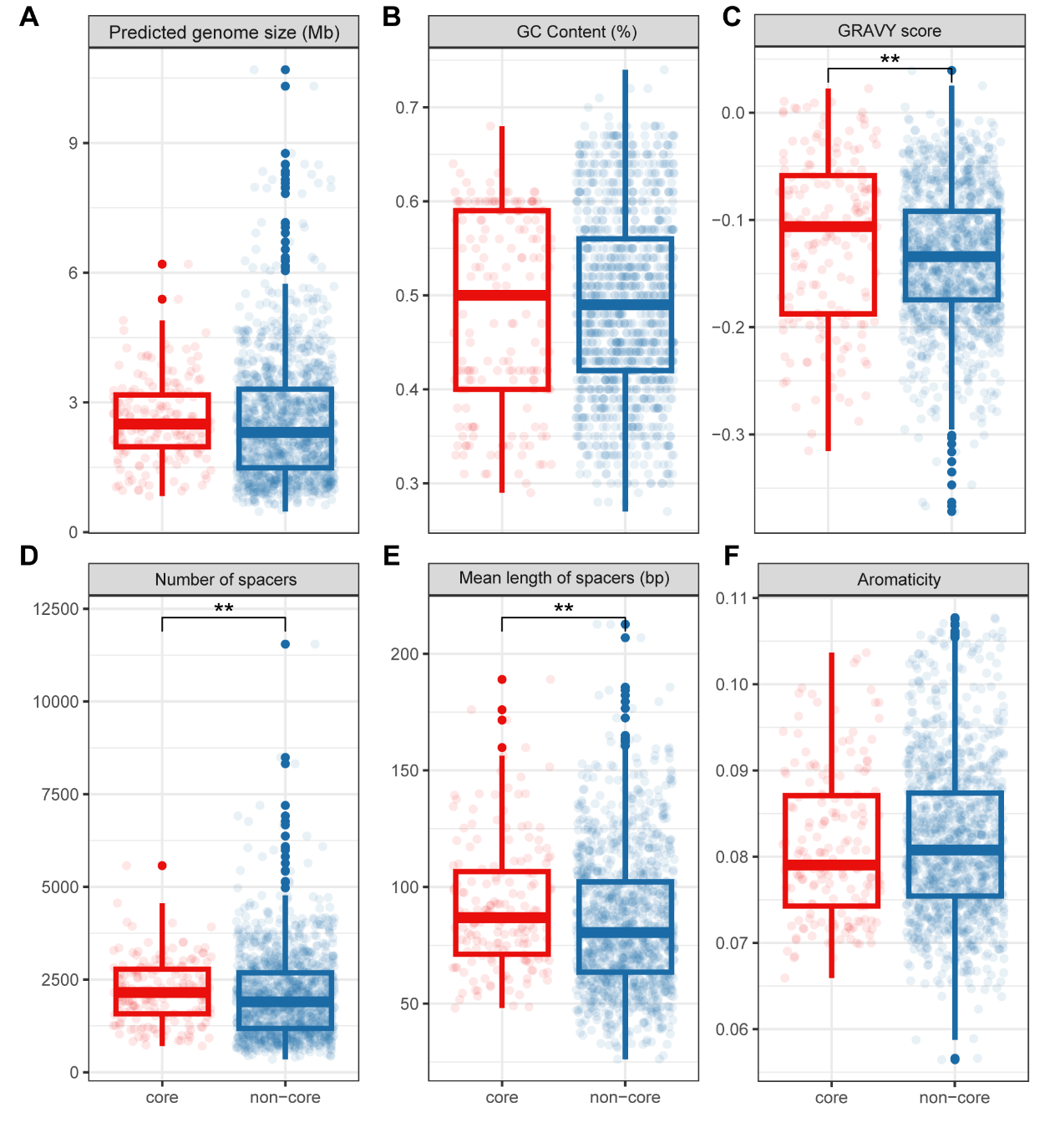

Figure S13. Comparative analysis of genomic traits among MAGs from core and non-core microorganisms at QDN-W07. (A-F) Genomic traits include (A) predicted genome size, (B) GC content, (C) grand average of hydropathy (GRAVY) score, (D) number of spacers, (E) mean length of spacers and (F) aromaticity. Red boxes and points represent MAGs of core microorganisms, while blue boxes and points represent non-core microorganisms. The significance level across groups was assessed using Wilcoxon rank-sum tests. Significance levels are denoted as: **, *p* ≤ 0.01. Source data are provided in Table S15.

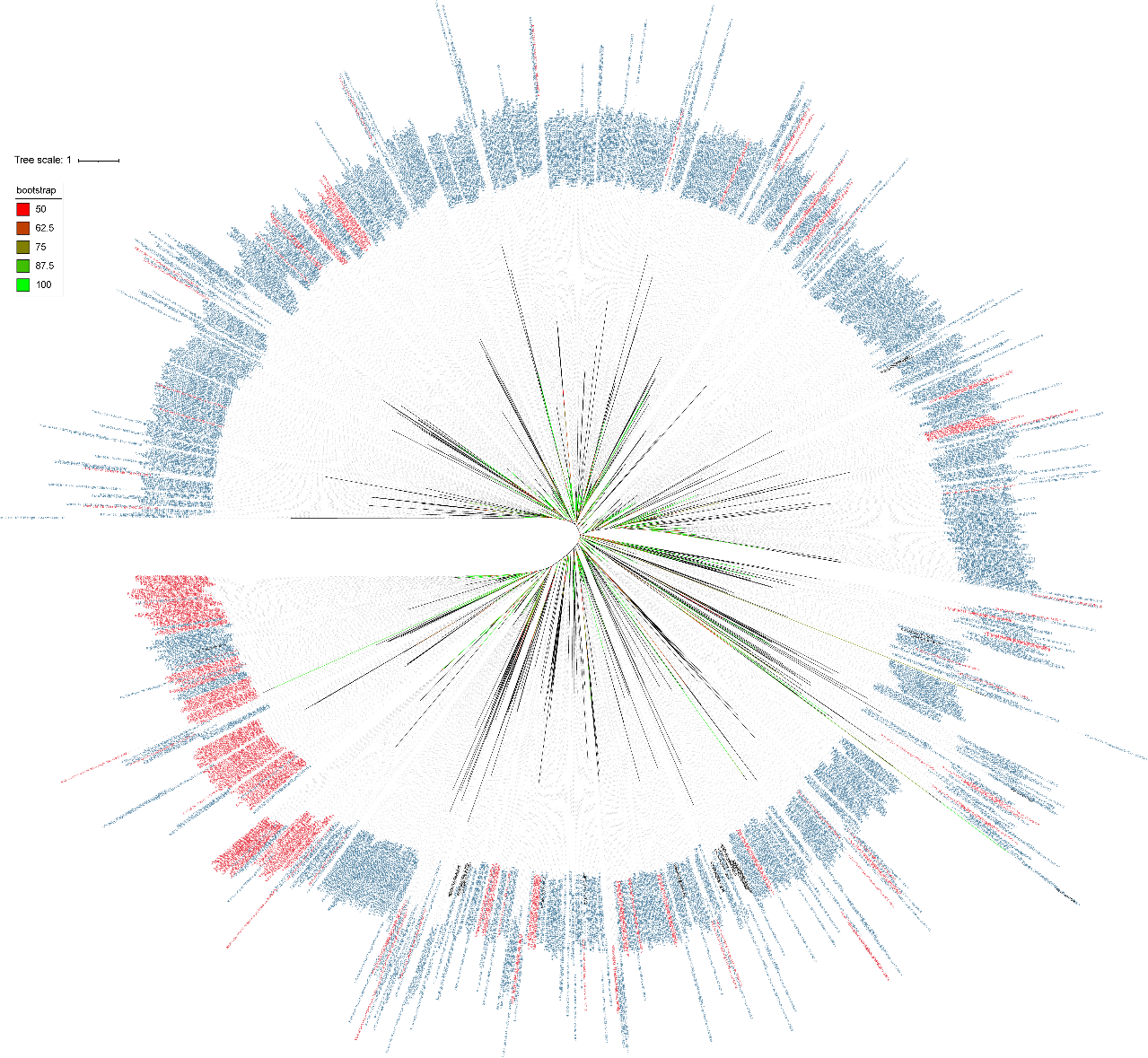

Figure S14. Gene phylogeny of cold shock proteins gene *cspA* identified from MAGs of QDN-W07. Nodes with red labels represent genes from core microorganisms, nodes with blue labels represent non-core microorganisms, and nodes with black labels represent reference sequences. Bootstrap values over 50% are indicated by different branch colors.

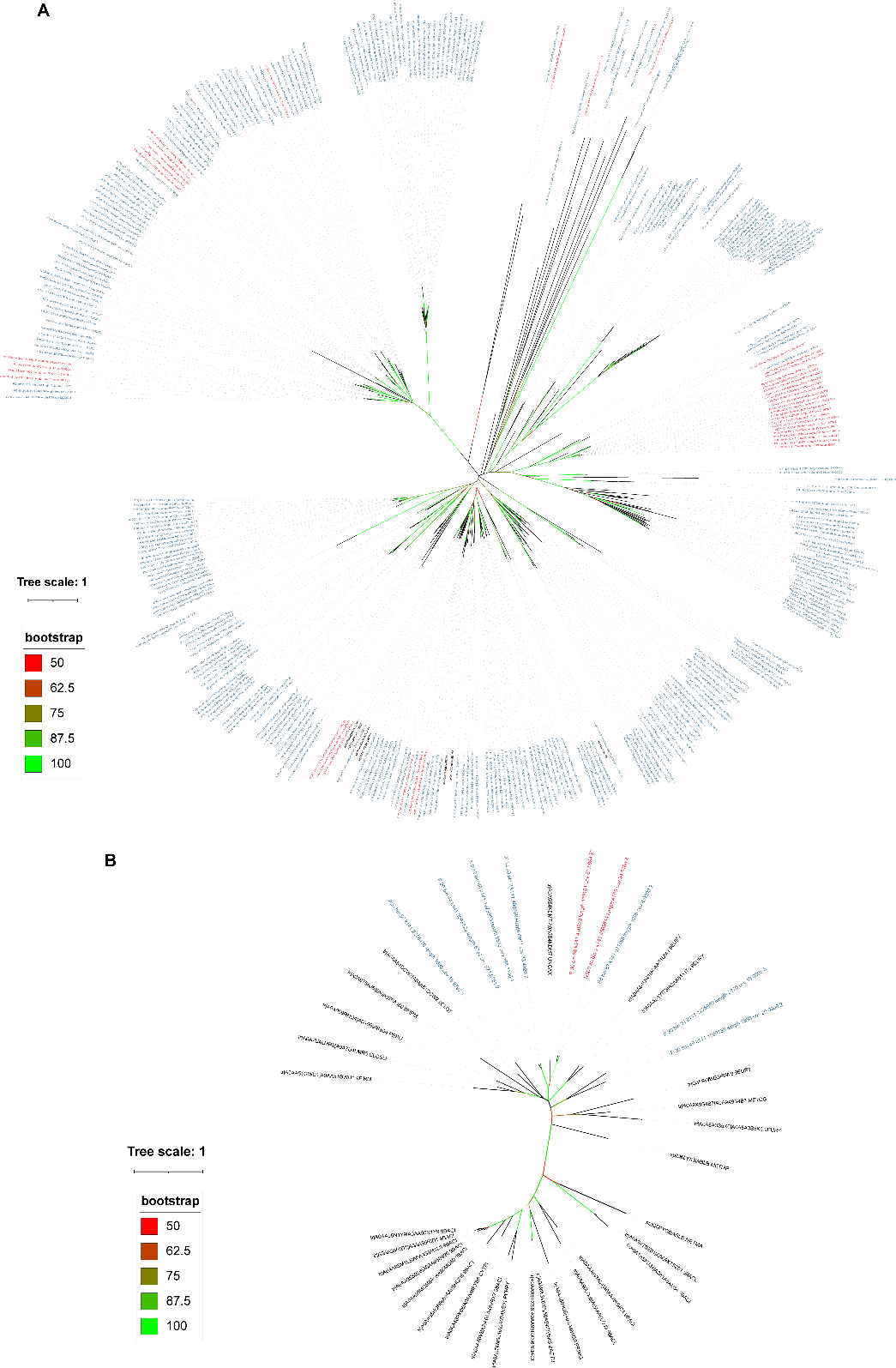

Figure S15. Gene phylogeny of salt adaptation genes identified from MAGs of QDN-W07. (A) *kamA* genes, which encode L-lysine 2,3-aminomutase, and (B) *ablB* genes, which encode Beta-lysine N(6)-acetyltransferase. Nodes with red labels represent genes from core microorganisms, blue labels represent non-core microorganisms, and black labels represent reference sequences. Bootstrap values over 50% are indicated by different branch colors.

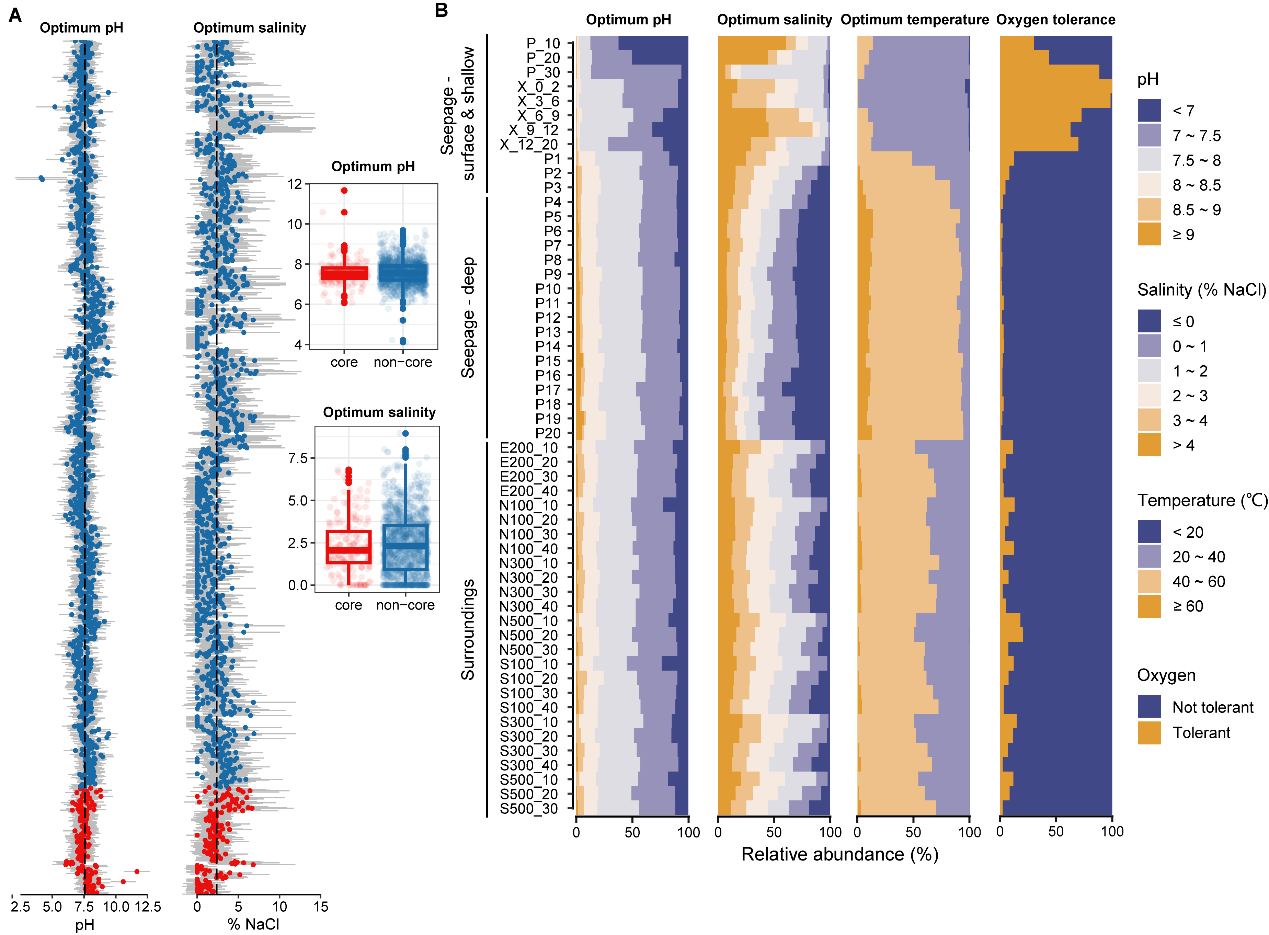

Figure S16. Predicted physicochemical growth conditions of microorganisms at QDN-W07. (A) Predicted physicochemical growth conditions (pH and salinity) of all MAGs, with mean values shown as dots and ±1 the root mean squared error (RSME) shown by lines. The vertical line indicates average values across all MAGs. Red points denote MAGs from core microorganisms, while blue points represent non-core microorganisms. Boxplots display comparisons of predicted pH and salinity between core and non-core microorganisms. (B) Predicted physicochemical growth conditions for microorganisms across different samples.

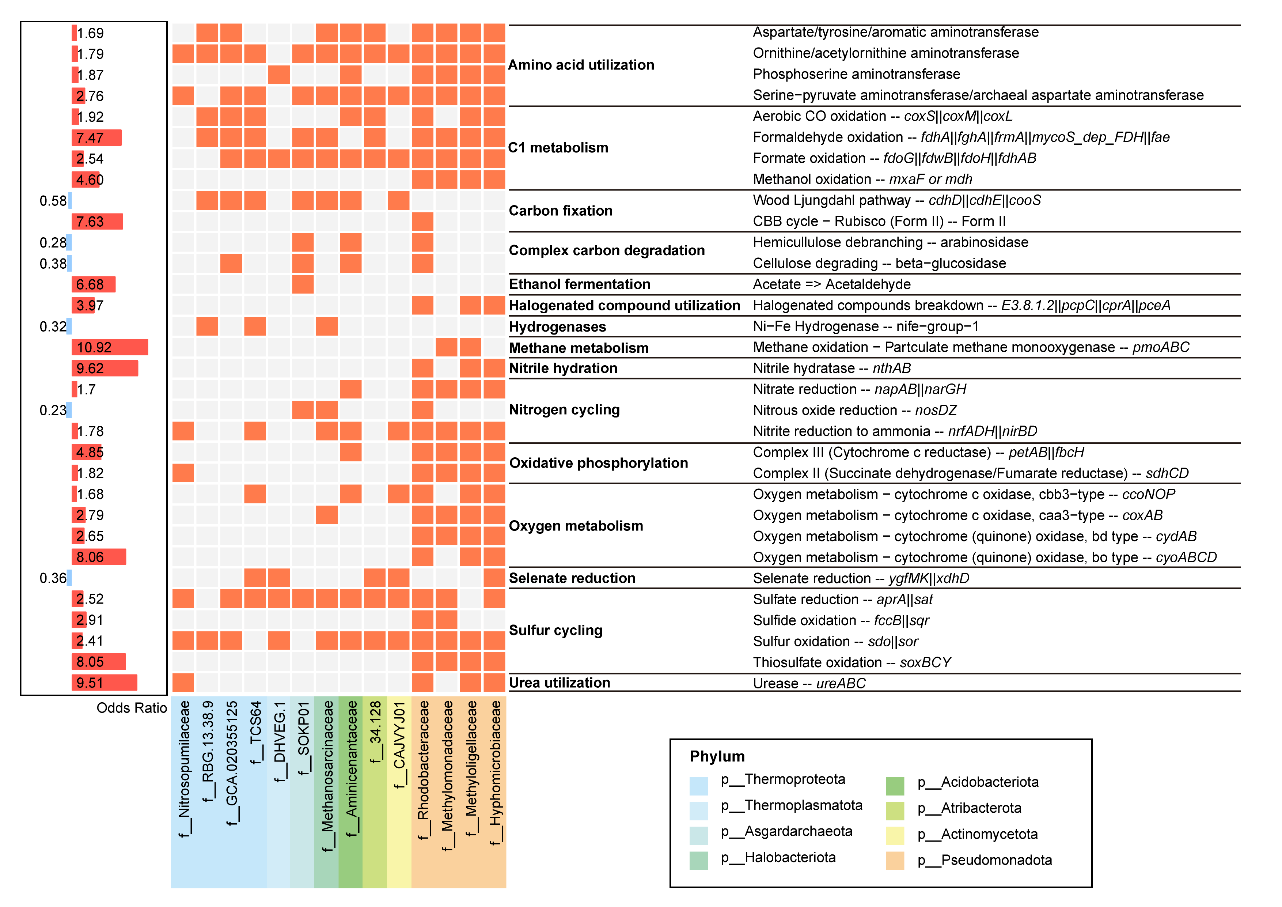

Figure S17. Functional enrichment in core microbiome. The chart displays the enrichment of metabolic functions within core microbiome, indicated by odds ratio (OR). Functions that are enriched in core microbiome are shown in red (OR > 1), while those that are not enriched are shown in blue (OR < 1). The heatmap displays the occurrence of these functions across specific microbial families, with orange indicating presence and grey indicating absence. Source data are provided in Table S17.

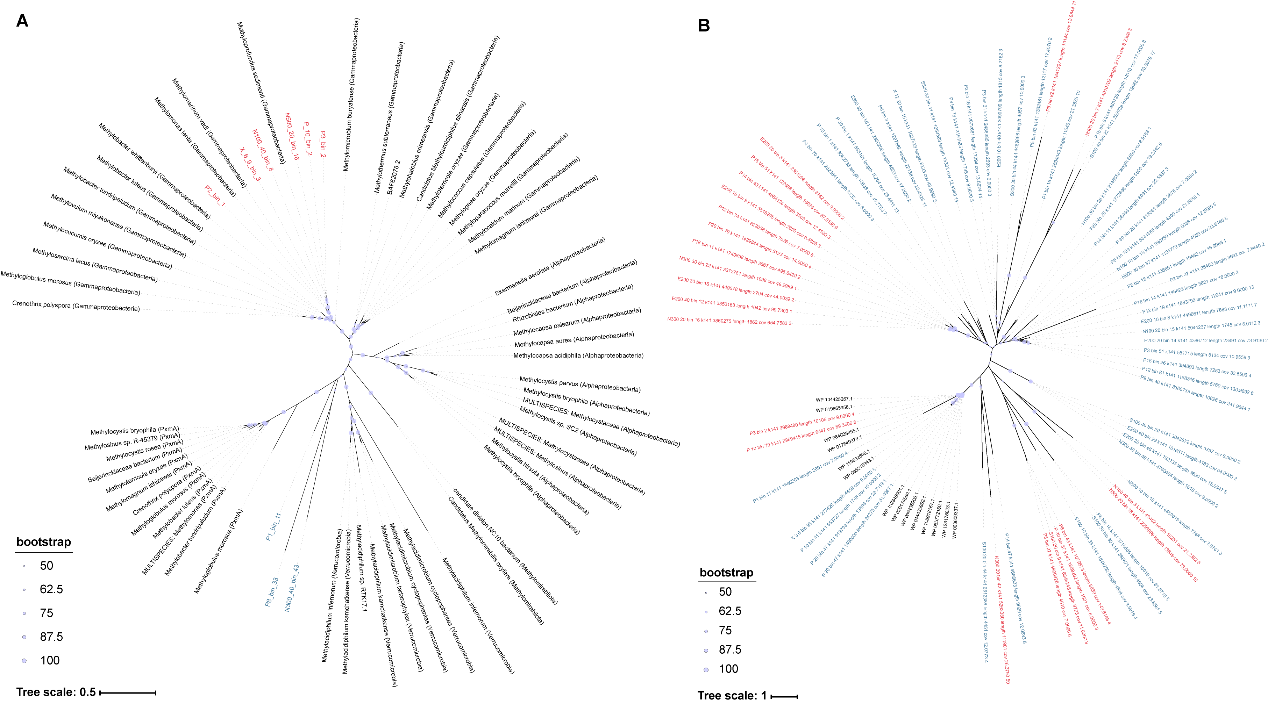

Figure S19. Gene phylogeny of aerobic methane oxidation genes identified from MAGs of QDN-W07. (A) *pmoA* genes, which encode α-subunit of particulate methane monooxygenase (pMMO), and (B) *mmoX* genes, which encode α-subunit of hydroxylase MMOH. Nodes with red labels represent genes from core microorganisms, blue labels represent non-core microorganisms, and black labels represent reference sequences. Bootstrap values over 50% are shown next to nodes.

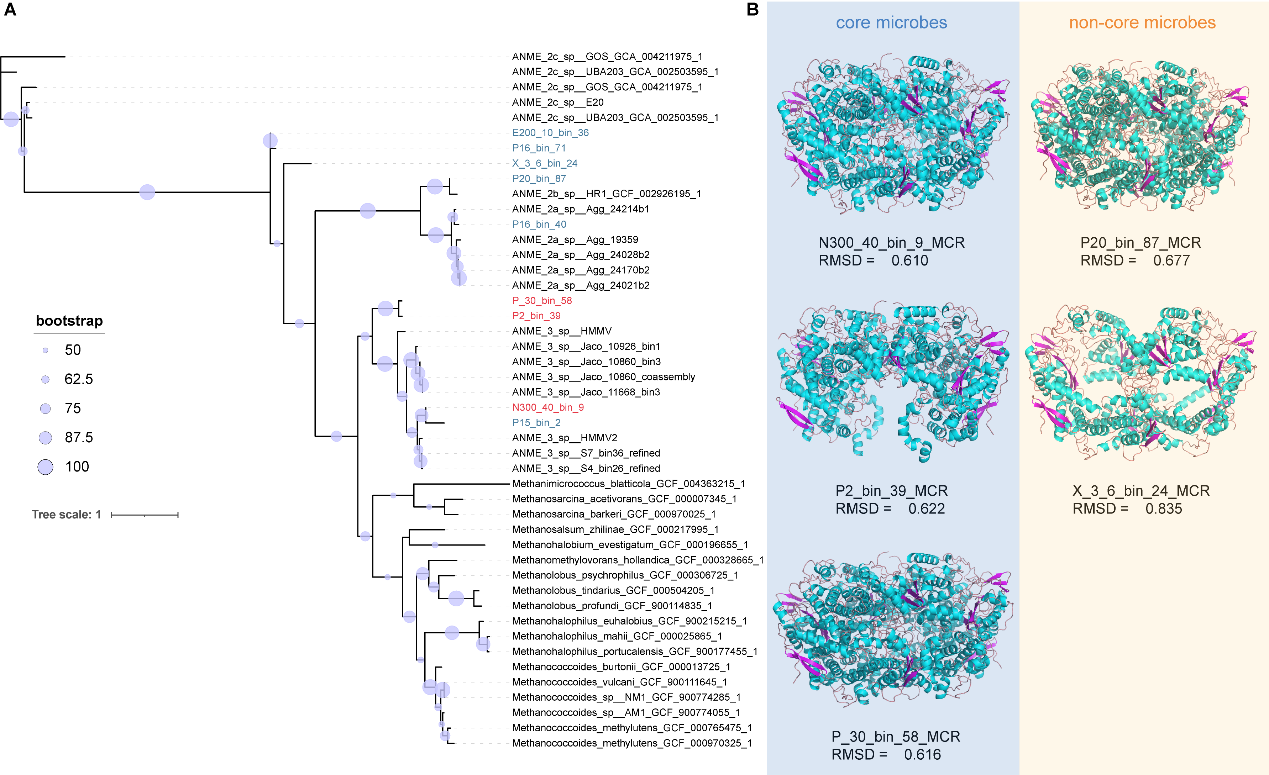

Figure S20. Phylogenetic analysis of *mcrA* genes and predicted structures of methyl-coenzyme M reductase (MCR). (A) Gene phylogeny of *mcrA* genes, which encode α-subunit of MCR. Nodes with red labels represent genes from core microorganisms, blue labels represent non-core microorganisms, and black labels represent reference sequences. Bootstrap values over 50% are shown next to nodes. (B) Predicted three-dimensional structures of MCR, including three proteins from core microbiome (with blue background) and two from non-core microbiome (with orange background). These structures were compared to the reference MCR structure from the PDB database (PDB ID: 1E6V). The comparison results are shown using Root Mean Square Deviation (RMSD) values, where RMSD < 2Å typically indicates a high degree of structural similarity between the two proteins.

Table S1. Sample information and microbial sequencing of three seep sites, QDN-W07, QDN-S18 and HM-ROV01, in the Haima cold seeps from the South China Sea.

Table S2. Geochemical data of the push cores and piston cores at QDN-W07, QDN-S18 and HM-ROV01 sites.

Table S3. Summary of ASV feature table at the phylum level.

Table S4. Relative abundance of key microbial taxa across various samples, including ANME, MOB and SRB.

Table S5. Network characteristics of microbial co-occurrence networks at QDN-W07, HM-ROV01 and QDN-S18 sites.

Table S6. Mantel test results correlating microbial communities with environmental factors.

Table S7. Relative contribution of different community assembly processes across groups.

Table S8. Mantel test results assessing the influence of homogeneous selection (HoS) and environmental factors.

Table S9. iCAMP results for archaeal communities of three seep sites, QDN-W07, QDN-S18 and HM-ROV01.

Table S10. iCAMP results for bacterial communities of three seep sites, QDN-W07, QDN-S18 and HM-ROV01.

Table S11. Core microbiome taxa identified from ASVs of QDN-W07 site.

Table S12. Core microbiome taxa identified from MAGs of QDN-W07 site.

Table S13. Relative abundance of MAGs of 54 samples of QDN-W07 site.

Table S14. Twenty amino acid count statistics in predicted proteins for 1365 MAGs.

Table S15. Genomic traits of 1365 MAGs.

Table S16. Gene enrichment in the core microbiome based on eggNOG-mapper analysis.

Table S17. Function enrichment in the core microbiome based on METABOLIC analysis.

Table S18. MAGs with key genes (*mcrA*, *mmoX* and *pmoA*) and their taxonomic classifications.
